# Novel enhancers of guanylyl cyclase-A activity via allosteric modulation

**DOI:** 10.1101/2021.12.31.474340

**Authors:** Henriette Andresen, Cristina Pérez-Ternero, Jerid Robinson, Deborah M Dickey, Adrian J Hobbs, Lincoln R Potter, Finn Olav Levy, Alessandro Cataliotti, Lise Román Moltzau

**Affiliations:** Department of Pharmacology, Division of Laboratory Medicine, University of Oslo and Oslo University Hospital, Oslo, Norway; William Harvey Research Institute, Barts & The London School of Medicine, Queen Mary University of London, London, United Kingdom; Department of Biochemistry, Molecular Biology, and Biophysics, University of Minnesota Medical School, Minneapolis, Minnesota, United States of America; Institute for Experimental Medical Research, University of Oslo and Oslo University Hospital, Oslo, Norway

## Abstract

Natriuretic peptide receptor (NPR)-A (also known as NPR-A, NPR1 or guanylyl cyclase-A, GC-A) is an attractive but challenging target to activate with small molecules. GC-A is activated by endogenous atrial natriuretic peptide (ANP) and B-type natriuretic peptide (BNP), and this activation leads to the production of 3’,5’-cyclic guanosine monophosphate (cGMP). This system plays an important role in the regulation of cardiovascular and renal homeostasis. However, utilization of this receptor as a drug target has so far been limited to peptides, even though small molecule modulators allow oral administration and longer half-life. We have identified small molecular allosteric enhancers of GC-A, which strengthened ANP or BNP activation in various *in vitro* and *ex vivo* systems. These compounds do not mediate their actions through previously described allosteric binding sites or via known mechanisms of action. In addition, their selectivity and activity are dependent on only one amino acid in GC-A. Our findings show that there is a novel allosteric binding site on GC-A, which can be targeted by small molecules that increase the signaling effects of ANP and BNP.

## INTRODUCTION

Natriuretic peptides (NPs) and their receptors are central regulators of cardiovascular and renal homeostasis, and enhancement of the natriuretic peptide system has become an attractive therapeutic target. Endogenous and several designer natriuretic peptides have been tested in the treatment of conditions such as heart failure^1–3^ and hypertension^4,5^, but the development of small molecular compounds has been challenging, and only a few have been developed to target this system^6–8^.

The NP system comprises three distinct hormones, two guanylyl cyclase-linked receptors and one clearance receptor with G-protein signaling. Atrial natriuretic peptide (ANP) and B-type natriuretic peptide (BNP) activate guanylyl cyclase (GC)-A (also known as NPR-A or NPR1) and C-type natriuretic peptide (CNP) activates GC-B (also known as NPR-B or NPR2). The activation of these two receptors produces the second messenger, 3’,5’-cyclic guanosine monophosphate (cGMP). In the heart and kidneys, GC-A activation has anti-remodelling effects as it inhibits hypertrophy and fibrosis. In the kidneys, GC-A activation also causes natriuresis and diuresis and enhances glomerular filtration rate. GC-A activation also inhibits the renin-angiotensin-aldosterone system (RAAS) in several tissues^9^. GC-A activation in the vasculature causes vasodilation, which, together with the renal actions and RAAS inhibition, reduces blood volume and pressure^10,11^. In animal models, genetic depletion of GC-A^12,13^, ANP^14, 15^ or BNP^16^ lead to hypertension, cardiac hypertrophy/fibrosis and organ damage, demonstrating the importance of GC-A activation in cardiovascular and renal homeostasis. In humans, genetic studies have demonstrated a link between altered GC-A function and blood pressure^17^ and the increased risk of development of hypertension^18^. Early corollary studies demonstrated that people with low NP concentrations had higher blood pressure than those with normal levels of NP^19,20^, which suggested the rationale for the development of therapeutic strategies aimed at enhancing this protective hormonal system.

GC-A is a homodimer with one NP binding site in the extracellular interface between the monomers^21^, which results in a ligand to receptor stoichiometry of 1:2. It is hypothesized that NP binding induces a small rotation of the two monomers that propagates through the single transmembrane spanning domain, the kinase homology domain (KHD), and the dimerization domain (CCD) to the GC catalytic domain. This leads to increased cGMP production as the two GC domains are brought closer together^22,23^. Besides the NP binding domain, several allosteric binding sites have been described on GC-A^24–30^. In addition to allosteric modulation, the activity of GC-A is affected by its phosphorylation state^31,32^ and post-translational glycosylation^33^.

Recombinant ANP and BNP have been used to target GC-A in the treatment of acute heart failure^34^. However, both these peptides also bind to NPR-C, which is known as a “clearance receptor” that internalizes and leads to lysosomal degradation of all NPs. Internalization through NPR-C and rapid enzymatic cleavage by neprilysin and other proteases are responsible for the short half-life of these peptides^35^. Therefore, researchers have pursued ways to increase the half-life by designing NPs that have increased resistance to neprilysin degradation^36,37^, or by inhibiting neprilysin. Valsartan/sacubitril is a combination of the angiotensin receptor blocker valsartan and the neprilysin inhibitor sacubitril and has been shown to be effective in the treatment of chronic heart failure^38^. Although the NP system is a validated therapeutic target, valsartan/sacubitril is currently the only small molecular compound on the market that modulates the NP system. This indicates that activation of GC-A with small molecules is challenging, and to our knowledge, only Iwaki et al. have successfully developed GC-A agonists that mimic ANP and BNP^6–8^. Small molecular compounds like these have, in contrast to peptides, the potential of oral administration and longer half-life.

Here, we describe allosteric enhancers of GC-A that are not competitive with ANP or BNP but require activation of GC-A for their *in vitro* and *ex vivo* effects. We have explored their mechanism of action and have discovered that they do not modulate the affinity of NPs, but increase their efficacy on GC-A. Further, by exploiting the selectivity of one of these compounds, we have identified a key amino acid that is necessary for the enhancing actions and we suggest a new allosteric binding site on GC-A that may be useful for further drug development.

## METHODS

### Cell cultures

QBIHEK293A HEK293T, and HeLa cells were grown in Dulbecco’s modified Eagle’s medium (DMEM) (Gibco®, ThermoFischer Scientific) that was supplemented with 10% fetal bovine serum (FBS), 100 U/ml penicillin and 0.1 mg/ml streptomycin. QBIHEK293A cells were stably transfected with human GC-A or GC-B, as previously described^39^. For the stable GC-A- and GC-B-expressing cells, 1 mg/ml geneticin (G418) was added as the selection antibiotic.

### The AlphaScreen assay for cGMP

The AlphaScreen assay for cGMP (PerkinElmer) was performed as previously described^39^. QBIHEK293 cells that expressed GC-A and GC-B were split the day before the experiment and harvested using an ethylenediaminetetraacetic acid (EDTA) solution (Versene, Invitrogen, ThermoFischer Scientific). Cells (6000/well GC-A-expressing, 8000/well GC-B-expressing) were resuspended in stimulation buffer (5 mM 4-(2-hydroxyethyl)-1-piperazineethanesulphonic acid (HEPES) in Hanks’ balanced salt solution (HBSS) at pH 7.4, 0.1% bovine serum albumin (BSA)) with isobutylmethyl xanthine (IBMX) (0.7 mM final). Compounds were dissolved in stimulation buffer and added to wells in the indicated concentrations. Cells were incubated with the indicated concentrations of compounds for 20 min before adding agonists (human BNP, CNP or proBNP) in various concentrations. Cells were stimulated for 20 min with agonist before the reactions were stopped/cells lysed by adding the AlphaScreen Acceptor bead mix (15.6 μg/ml AlphaScreen Protein A-coated Acceptor Beads (final concentration 3.13 μg/ml), anti-cGMP antibody (PerkinElmer anti-cGMP antibody: final dilution1:8000, Genscript anti-cGMP antibody: final dilution 1:50,000) and 0.5% Tween-20 in 5 mM HEPES buffer at pH 7.4). After incubation for 1h, the Donor bead mix (7.8 μg/ml AlphaScreen Streptavidin Donor beads (final concentration 3.13 μg/ml), biotinylated cGMP (PerkinElmer biotinylated cGMP: 0.625 nM final concentration, BIOLOG biotinylated cGMP: 6.25 nM final concentration), 0.5% Tween-20 in 5 mM HEPES buffer at pH 7.4) was added (40 μl final volume) and incubation continued for 2 h. The luminescence signals were quantified on an EnVision® multilabel plate reader (PerkinElmer) using AlphaScreen emission 570 nm filter.

When transiently transfected cells were used, QBIHEK293A cells were grown to 70-80% confluency and transfected using Lipofectamine LTX Plus (Invitrogen) according to the manufacturer’s instructions or by use of polyethyleneimine (PEI) in a 3:1 PEI:DNA ratio. After 48 h, cells (16,000/well) were harvested using TrypLE™ Select Enzyme (Gibco®, ThermoFisher Scientific) and the assay was carried out as described above.

All test compounds were dissolved in dimethyl sulphoxide (DMSO), the concentration of which was kept constant or included in controls for all experiments. Data were analyzed using GraphPad Prism 8.3.0 software. For the construction of concentration-response curves for NPs, curves were fitted using nonlinear regression and the built-in log (agonist) vs. response (three parameters) (Hill slope = 1). To construct concentration-response curves for the compounds in the presence of a small concentration of NP, the curves were fitted using log (agonist) vs. response – variable slope (four parameters).

### High throughput screening

Chemical libraries from Enamine (28,500 compounds) and a protein-protein interaction library (1,008 compounds) were screened using the AlphaScreen assay for cGMP. Compounds (10 μM final concentration in 40 μl) were printed on the plates by a Labcyte Echo 550 Acoustic Liquid Handler. Cells (8000/well) were added by a Hamilton Microlab Star automated liquid-handling robot and incubated for 20 min before they were stimulated with a small concentration of rat BNP (EC_10_; 3 nM). This concentration was chosen to enable the identification of both agonists and allosteric modulators. Maximum cGMP production was induced by the addition of 300 nM rat BNP as a control to every plate. Reagents were added with PerkinElmer FlexDrop PLUS to a total volume of 40 μl. Plates were read on a PerkinElmer EnVision® multilabel plate reader using Turbo option.

### Membrane preparation and binding assay

#### Membrane preparation

GC-A-expressing QBIHEK293A cells were harvested in ice-cold HBSS and collected by centrifugation (800 × g, 5 min, 4 °C). The pellet was resuspended in ice-cold homogenization buffer (STE: 27% (w/w) sucrose, 50 mM Tris-HCl, pH 7.5 at RT, 5 mM EDTA) and homogenized with an Ultra-Turrax homogenizer. The homogenate was pelleted at 300 × g for 5 min at 4 °C, and the supernatant was centrifuged at 27,000 × g for 20 min at 4 °C. The pellet was resuspended in ice-cold 50 mM Tris-HCl, pH 7.5 at RT, and 1 mM EDTA using a Dounce glass-glass homogenizer. It was centrifuged at 27,000 × g for 20 min at 4 °C, and the pellet was resuspended again and homogenized.

#### Binding assays using membranes

Binding reactions were performed in a 50 μl volume that contained 0.5 μg of membranes, binding buffer (50 mM Tris-HCl, pH 7.5 at RT, and 0.1 mM EDTA, 5 mM MgCl_2_ and 0.1% BSA), 50 pM ^125^I-ANP, and the indicated concentrations of NP and/or compounds or DMSO in a 96-well format. Competition binding in the presence of 1 mM adenosine triphosphate (ATP) was performed similarly, but with the addition of an ATP-regenerating system to all wells (20 mM creatine phosphate, 0.2 mg/ml creatine phosphokinase, 40 U/ml myokinase final). The reactions were incubated for 2.5 h at RT before membranes were harvested onto Millipore harvest plates that had been pre-soaked in 1% PEI through the use of a Packard Cell Harvester. Then the membranes were washed four times with cold 50 mM potassium phosphate buffer (pH 7.4 at RT). The filter plates were dried and 20 μl MicroScint™-O cocktail scintillation fluid (PerkinElmer) were added to each well before the plate was counted in a Packard TopCount Scintillation Counter (Packard Instrument Co.).

#### Whole cell binding assays

This protocol was adapted from Dickey et al.^40^. QBIHEK293 cells that expressed GC-A were seeded equally into 24-well plates that had been coated with poly-L-lysine and grown to 90% confluency. The following day, cells were pretreated with DMEM that had been supplemented with 0.2% BSA for 1-2 h at 37 °C. For the competition binding assays, cells were incubated in binding buffer (DMEM supplemented with 1% BSA) with 50 pM ^125^I-ANP and the indicated concentrations of NP and/or compounds or DMSO. For the saturation binding assays, cells were incubated in the same binding buffer with varying concentrations of ^125^I-ANP and 0.1% DMSO (control) or 10 μM of the compound. For nonspecific binding, 1 μM ANP was added to the cells. Cells were incubated at 4 °C for 1 h. After incubation, cells were washed twice with ice-cold phosphate-buffered saline (PBS) (pH 7.4 at RT) and the cells were harvested in 500 μl 1 M NaOH. The solution was transferred to scintillation vials and radioactivity was counted using Ultima Gold XR scintillation cocktail (PerkinElmer) in a liquid scintillation counter (Tri-Carb 2300 TR, Packard).

### Phosphodiesterase activity assay

Cells were homogenized in ice-cold 20 mM Tris-HCl (pH 7.5 at RT) that contained 1 mM EDTA, 1 mM dithiotreithol, 0.2 mM phenylmethylsulphonyl fluoride (PMSF), 0.1 mM sodium orthovanadate (Na_3_VO_4_), 1 mM benzamidine, 20 μg/ml leupeptin and 10 μg/ml aprotinin (trypsin inhibitor). The cGMP phosphodiesterase (PDE) activity was then measured by a modified two-step procedure^41^, with modifications as previously described^42^.

### Substrate-velocity assay

GC assays were performed at 37 °C in a reaction mixture that contained 25 mM HEPES (pH 7.4), 50 mM NaCl, 0.1% BSA, 0.5 mM IBMX, 1 mM EDTA, 5 mM phosphocreatine, creatine kinase (0.1 mg/ml), 0.5 mM microcystin, and 5 mM MgCl_2_. Guanosine triphosphate (GTP) concentrations were as indicated in the figures. Reactions were initiated by adding 20 ml of crude membranes from GC-A-expressing HEK293T cells that contained 10-18 mg of protein suspended in phosphatase inhibitor buffer to 80 ml of the reaction mixture. Reactions were terminated in 110 mM ZnOAc and 110 mM Na_2_CO_3_ and purified on acidified alumina after elution in 3 ml of 200 mM ammonium formate to separate ATP and GTP from cGMP^32^. Concentrations of cGMP were determined by the performance of enzyme-linked immunosorbent assays (ELISA) according to the manufacturer’s instructions (ENZO Lifesciences).

### Isolation of rat aortic rings and measurements of vascular reactivity

All animal studies followed the UK Animals (Scientific Procedures) Act of 1986 and had approval from a local Animal Welfare and Ethical Review Body. Male Wistar rats (six to eight weeks old) were killed by CO_2_ asphyxiation. The thoracic aorta of each was dissected and rings (~4 mm length) were mounted in organ baths that contained physiological salt solution (PSS: 119 mM NaCl, 4.7 mM KCl, 2.5 mM CaCl_2_, 1.2 mM MgSO_4_, 25 mM NaHCO_3_, 1.2 mM KH_2_PO_4_, and 5.5 mM glucose), which was maintained at 37 °C and gassed with 5% CO_2_ in O_2_. Changes in isometric tension were measured in the tissues under a basal tension of 1 g. The viability of the tissue was assessed by exposure to KCl (80 mM). Then, the maximal contractile response was recorded by exposure to a single dose of the thromboxane receptor agonist 9, 11-dideoxy-11α, 9α-epoxymethano-prostaglandin F2α (U46619; 1 μM). Arteries were then treated with the nitric oxide synthase (NOS) inhibitor L-N^G^-nitroarginine methyl ester (300 μM) to block the production of endogenous nitric oxide. The arteries were then pre-contracted with an 80% maximal effective (EC_80_) concentration of U46619. Once a stable response had been achieved, cumulative concentration–response curves were constructed either with increasing concentrations of compounds #2 or #20, or with ANP in the absence or presence of 10 μM of compound #2 or compound #20. In the latter case, compounds were incubated for 30 min prior to the administration of ANP.

### Cell-based cAMP assay

The level of NPR-C agonism was assessed by quantifying the inhibition of forskolin-induced cyclic adenosine-3’,5’-monophosphate (cAMP) production in HeLa cells. Cells were grown to around 80% confluency in 12-well plates and compounds (30 μM) or the specific NPR-C agonist cANF^4-23^ (100 nM) were added and left to equilibrate with the cells for 10 min before the addition of forskolin (10 μM). In some cases, the NPR-C antagonist osteocrin (100 nM) was added 10 min before the compounds were added. The reaction was stopped 20 min after forskolin addition and cells were lysed in HCl (0.1 M) and centrifuged (18,620 × g, 2 min, 4 °C). Concentrations of cAMP were measured in the supernatants through use of an ELISA (Direct cAMP; Enzo Life Sciences) following the manufacturer’s instructions and the values were normalized according to the protein concentration.

### Construction of chimeric GC-A/B and mutated receptors

Site-directed mutagenesis in GC-A and GC-B and construction of chimeric receptors were performed through use of the In-Fusion HD plus cloning kit (Takara Bio Inc.). The primers used are listed in Supplementary Table 1. All plasmid sequences were verified by Sanger sequencing (Eurofins Genomics, GmbH).

### Measurement of levels of cGMP in isolated rat cardiac fibroblasts

Cardiac fibroblasts were isolated from Wistar rat hearts. Animal care was conducted according to the Norwegian Animal Welfare Act (and approved by the Norwegian Animal Research Authority), which conforms with the Directive 2010/63/EU of the European Parliament and of the European Council of 22 September 2010 on the protection of animals used for scientific purposes. Hearts were perfused at 37 °C using a Langendorff set-up with a buffer (Buffer A: 24 mM NaHCO_3_, 0.6 mM MgSO_4_, 1 mM DL-carnitine, 10 mM creatine, 20 mM taurine and 0.1% BSA) in Joklik-modified minimum essential medium (MEM) (one ampoule/l Joklik-MEM; M0518, Sigma-Aldrich) at a constant 6.4 ml/min flow of equilibrated 5% CO_2_/95% O_2_. Collagenase Type-II (90 U/ml final) (Worthington Biochemical Corporation, 268 U/mg) was added after 6 min and CaCl_2_ (0.2 mM final) after 24 min. When the aortic valves had been digested (33-35 min), the ventricles were minced and gently shaken at 37°C for 10 min in Buffer A with collagenase Type-II and 0.2 mM CaCl_2_ with continuous 5% CO_2_/95% O_2_ supply. The suspension was filtered (nylon mesh, 250 μm) and centrifuged at 30 × g for 3 min at RT. The supernatant was kept (Supernatant 1) and the pellet was resuspended in 15-20 ml of buffer B (Buffer A plus 1% BSA and 0.5 mM CaCl_2_). The suspension was centrifuged once more at 30 × g for 4 min at room temperature and the supernatant was combined with Supernatant 1. The combined supernatant was centrifuged at 1000 × g for 5 min at room temperature. The cell pellet was resuspended in Buffer A plus 0.5 mM CaCl_2_, 1% penicillin-streptomycin and 10% FBS to appropriate volume and incubated at 37 °C for 2 h before the medium was changed. The fibroblasts were grown in DMEM (Gibco®, ThermoFischer Scientific) supplemented with 10% FBS, 100 U/ml penicillin, 0.1 mg/ml streptomycin and 0.25 mg/ml amphotericin B and plated to 6-well plates in P2. The medium was changed to a serum-free medium 24 h before experiments were conducted. Cells were pre-incubated with compound #20 for 20 min and stimulated for 20 min with agonist as indicated. The experiment was stopped by the addition of 5% trichloroacetic acid (TCA) and the concentration of cGMP was measured using the cGMP enzyme immuno assay (EIA) kit (Cayman Chemical Company).

### Materials

^125^I-ANP was purchased from Phoenix Pharmaceuticals. NPs were obtained from GenScript and Sigma-Aldrich and proBNP from HyTest. Anti-cGMP antibody was obtained from PerkinElmer and GenScript. Biotinylated cGMP was bought from PerkinElmer and BIOLOG. Compounds #2 and #20 were obtained from MolPort and Mercachem. Compound #20 was also synthesized by Drug Discovery Laboratory, AS.

## RESULTS

### Identification of small molecular GC-A modulators

To identify activators of GC-A, we performed a high throughput screening. The assay was evaluated and validated based on Z’ values^43^. Z’ values were considered satisfactory for a cell-based assay (0.44±0.19). The cut-off for hit compounds was set to >30% stimulation, and these compounds were rescreened in triplicate and counter-screened using non-transfected QBIHEK293 cells. About 100 compounds were identified as hits and rescreened in triplicate. Only compound #2 (Fig. 1) was verified as a hit. We tested over 100 analogues of compound #2 for activity towards GC-A in a hit-to-lead process. Compound #20 was identified as the most potent compound; it had an almost 10-fold higher potency than compound #2 (EC_50_ values of 508±67 nM (#20) vs. 3,300±800 nM (#2)) (Fig. 2a). The concentration-response curves for compounds were obtained in the presence of 0.1 nM human BNP, which generated 10% of maximum cGMP production (EC_10_). Compounds #2 and #20 showed similar efficacies and increased the cGMP production to 45%±5% and 35%±3% of the maximum BNP-mediated cGMP production, respectively.

**Fig. 1.**
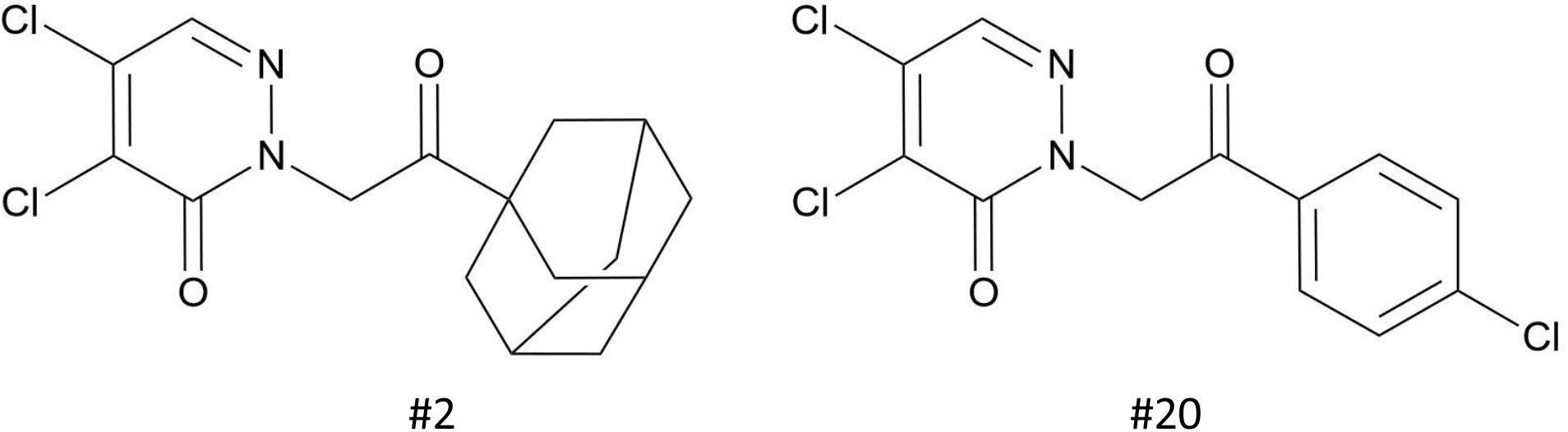
Molecular structures of compound #2, IUPAC name: 2-[2-(1-adamantyl)-2-oxoethyl]-4,5-dichloropyridazin-3-one;and of compound #20, IUPAC name: 4,5-dichloro-2-[2-(4-chlorophenyl)-2-oxoethyl]-2,3-dihydropyridazin-3-one.

**Fig. 2.**
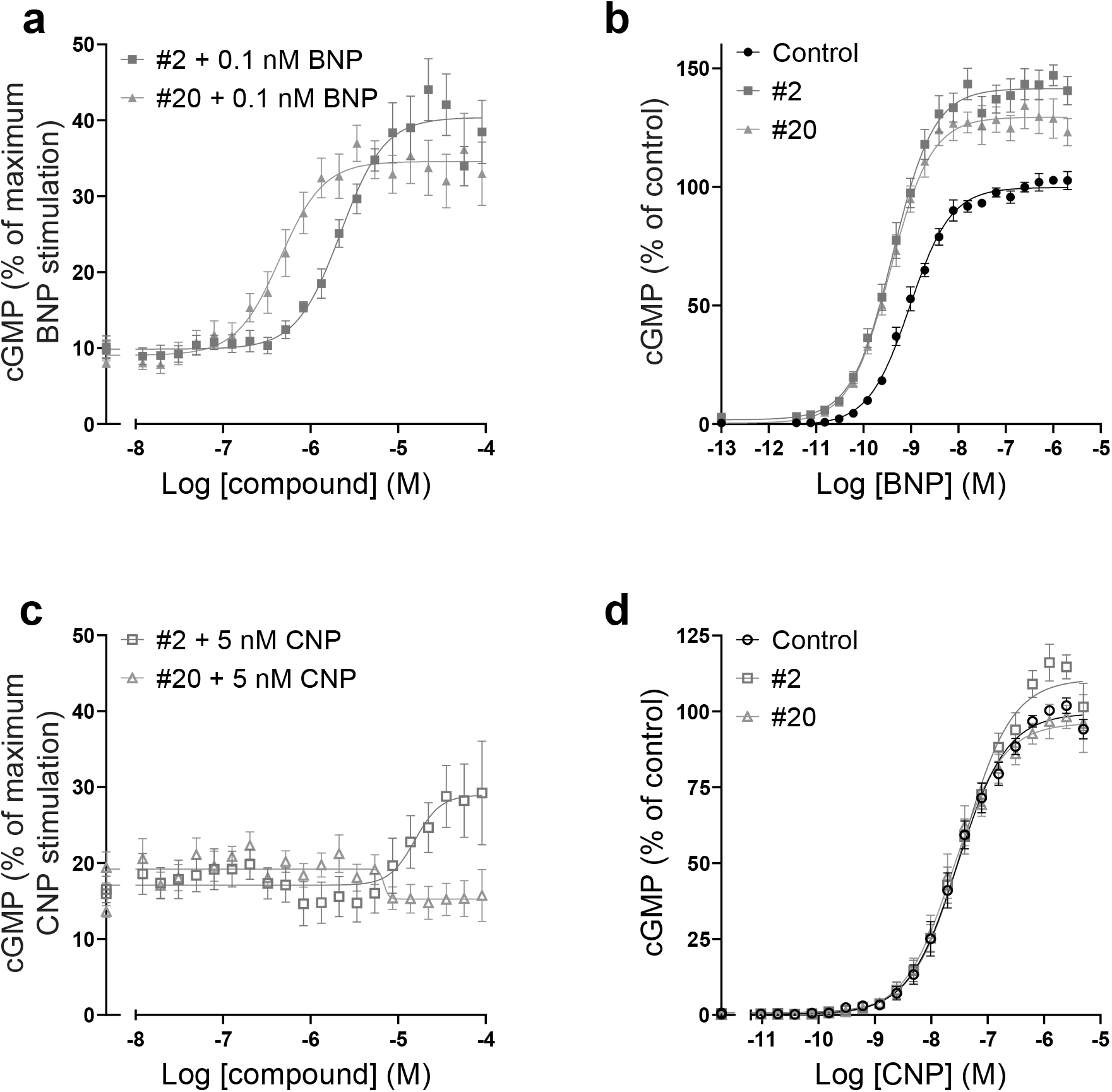
Compounds #2 and #20 enhanced BNP-induced GC-A, but not CNP-induced GC-B activity. **a**, **c**, Concentration-response curves for compounds #2 and #20 in the presence of an EC_10_ of NP as indicated. The production of cGMP was normalized to the maximum NP-mediated cGMP production in each experiment (n=5-7). Concentration-response curves for BNP (**b**) and CNP (**d**) in the absence or presence of 10 μM of compound #2 or #20 in cells that expressed GC-A and GC-B, respectively (n=6). Data points are means±SEM.

None of the compounds increased cGMP production in the absence of BNP, but both compounds #2 and #20 enhanced the effects of BNP (Fig. 2b). These two compounds increased the potency of BNP (EC_50_: 1±0.4 nM) by 2.4±0.5-fold (#2) and 2.8±0.4-fold (#20) and increased the maximum BNP-mediated cGMP production by 42%±5% and 30%±6%, respectively.

### Compound #20 showed selectivity towards GC-A

To determine selectivity towards GC-A, compounds were tested for activity towards GC-B and NPR-C. Compound #2 showed some activity towards GC-B at high concentrations, while compound #20 did not (Fig. 2c). Neither of the compounds modulated the potency of CNP or increased the maximum level of CNP-mediated cGMP production (Fig. 2d).

Compounds were tested towards NPR-C by measuring NPR-C-induced inhibition of cAMP (Fig. 3). The presence of compound #20 did not reduce cAMP production and thus did not activate NPR-C, while the presence of compound #2 reduced cAMP production and thus activated NPR-C. This reduction was reversed when the NPR-C antagonist osteocrin was added.

**Fig. 3.**
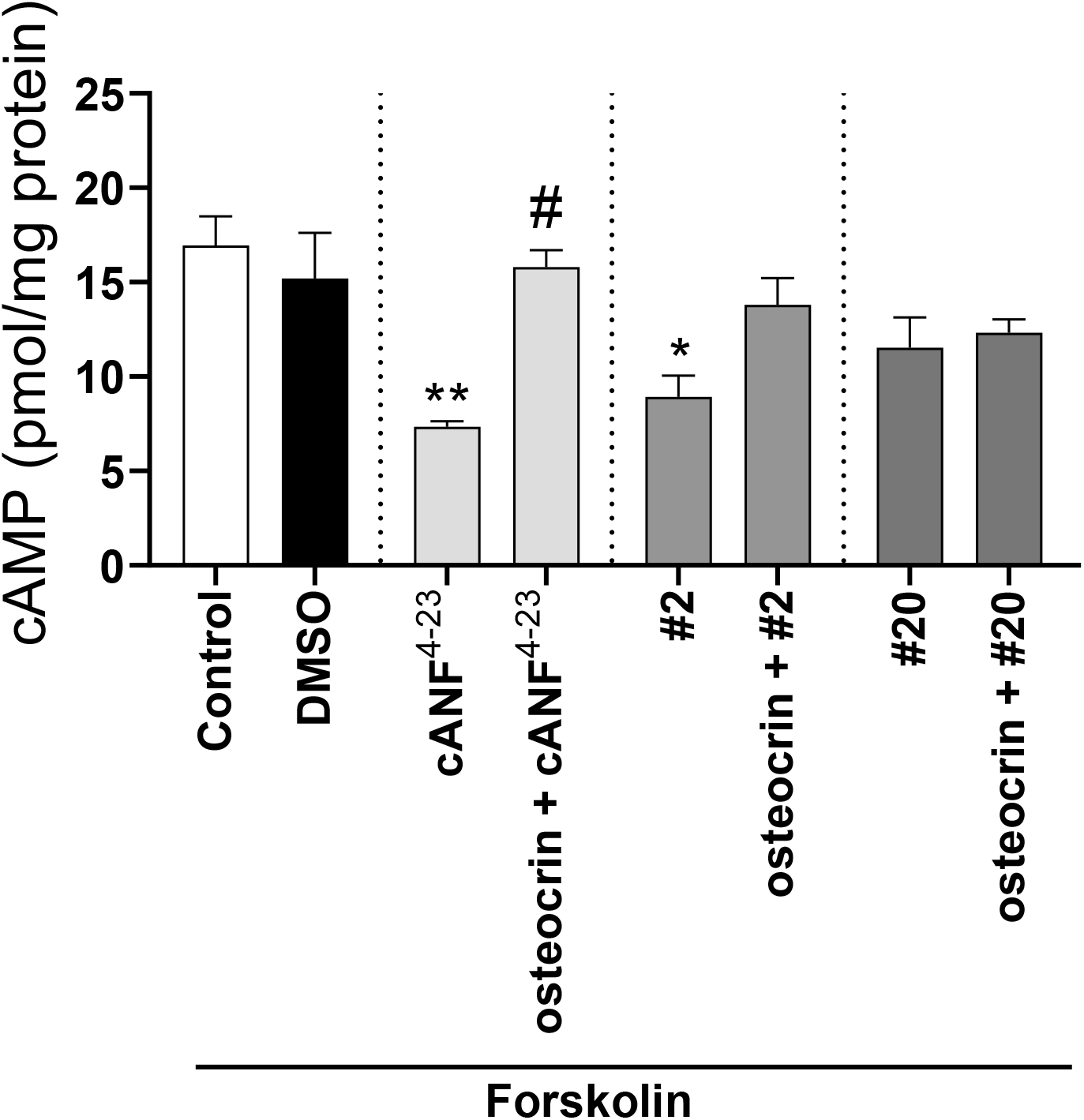
Compound #20 did not activate NPR-C. Forskolin-mediated cAMP production was inhibited by NPR-C activation in HeLa cells. Addition of the selective NPR-C agonist cANF^4-23^ (100 nM) as a positive control reduced cAMP accumulation and this effect was reversed by the addition of the NPR-C antagonist osteocrin (100 nM) prior to stimulation. To investigate whether our compounds modulated NPR-C activity, 30 μM of compounds were added prior to forskolin (10 μM) stimulation and osteocrin was added to investigate whether this reversed the effect. Data points are means±SEM (n=5). **P<0.01, *P<0.05 vs. forskolin, #P<0.05 v cANF^4-23^, one-way ANOVA.

To rule out the possibility that the enhancement of GC-A-induced production of cGMP by our compounds was due to effects on PDEs, cGMP-PDE activity was measured in homogenates from GC-A-expressing cells in the presence of our compounds. Neither of our compounds inhibited PDE activity alone or in the presence of IBMX (Supplementary Fig. S1).

### Compounds modulated the effects of unprocessed BNP

Secretion of unprocessed BNP is increased in heart failure and may be of clinical significance^44^. The BNP-precursor, proBNP, activates GC-A with lower potency than BNP^45^. We investigated whether compound #2 or #20 could modulate proBNP-mediated production of cGMP. When increasing concentrations of proBNP were co-incubated with 10 μM of compounds, compounds #2 and #20 increased cGMP production by 43%±10% and 42%±9%, respectively, but did not change the EC_50_ for proBNP (Fig. 4a). In the presence of a low concentration of proBNP, compound #20 increased cGMP production in a concentration-dependent manner by 277%±41%, and had an EC_50_ of 1.1±0.2μM (Fig. 4b).

**Fig. 4.**
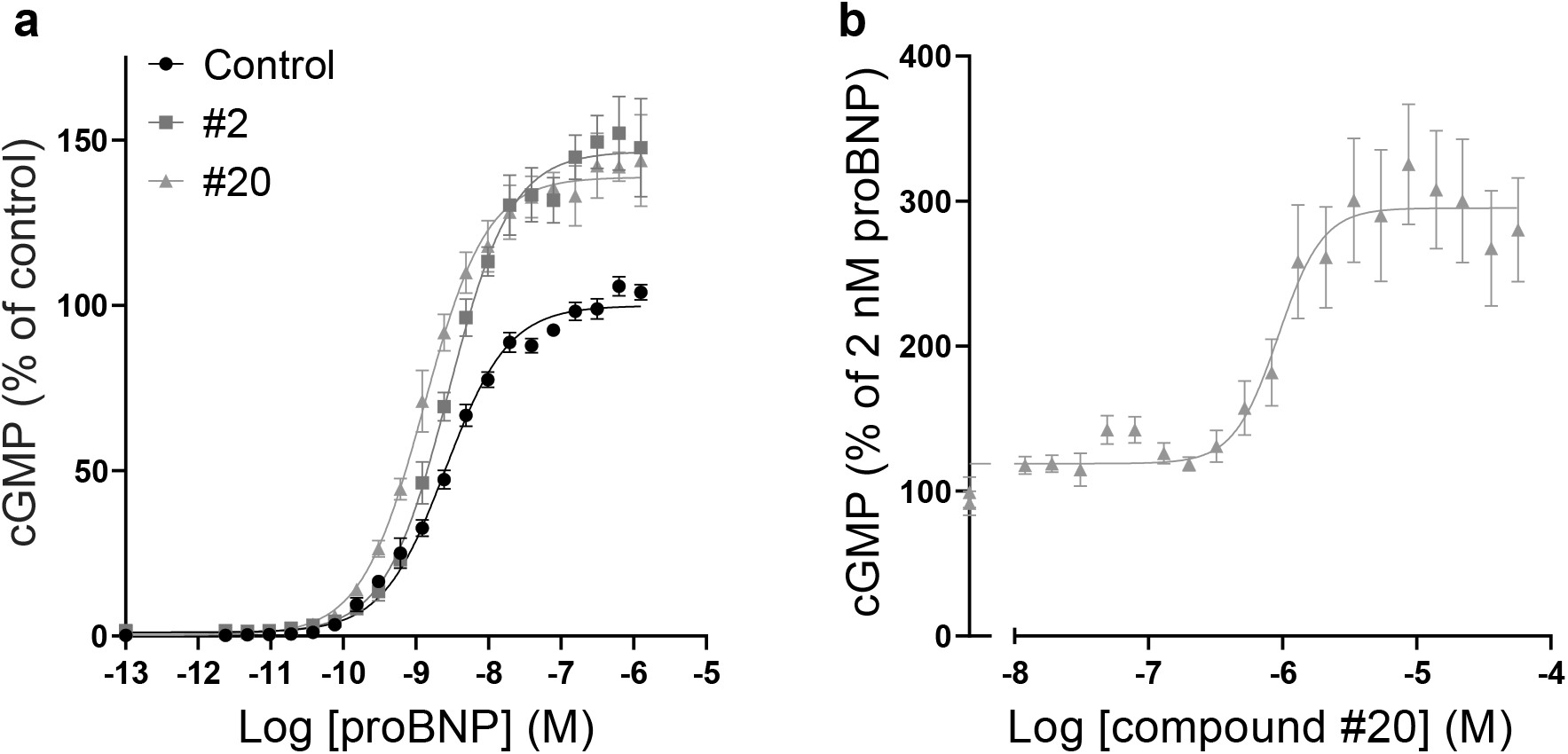
Compounds enhanced unprocessed BNP stimulation of GC-A. **a,** Concentration-response curves for proBNP in absence or presence of 10 μM compound #2 or #20 in GC-A-expressing cells. **b,** Concentration-response curve for compound #20 in the presence of 2 nM proBNP. The production of cGMP was normalized to proBNP alone (100%). Data points are means±SEM (n=4-6).

### Compound #20 increased BNP-mediated cGMP production in cardiac fibroblasts

In contrast with cell lines that overexpressed GC-A, cardiac fibroblasts endogenously express GC-A and activation inhibits cardiac fibrosis^9^. We wanted to investigate the effects of our compounds in a system that contained physiological expression levels of GC-A and stimulated isolated rat cardiac fibroblasts with BNP in the presence of DMSO or compound #20. The presence of compound #20 alone did not increase cGMP concentrations, but doubled the cGMP production (2.2±1.2-fold) in the presence of BNP compared with BNP alone (Fig. 5).

**Fig. 5.**
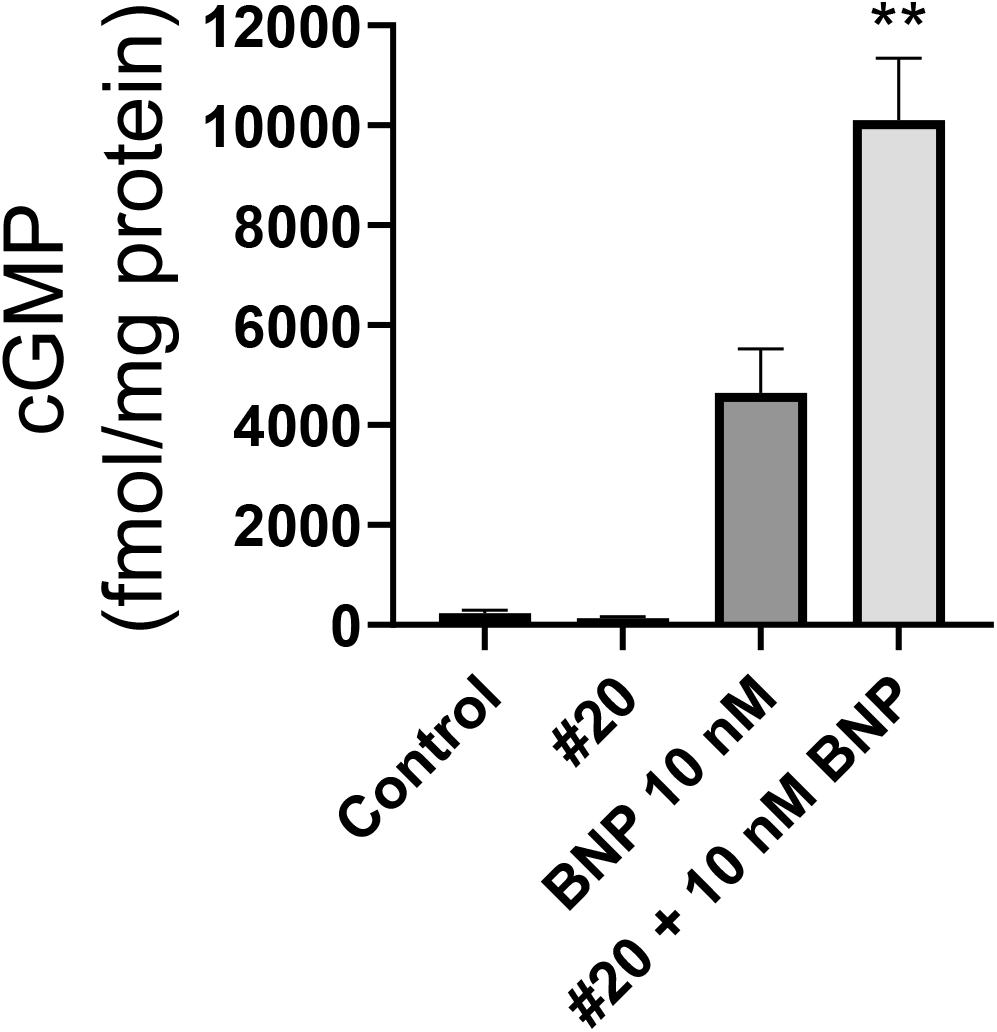
Compound #20 increased levels of BNP-mediated cGMP in rat cardiac fibroblasts. Levels of cGMP in rat cardiac fibroblasts at basal levels (control), 10 μM #20, 10 nM BNP and pretreatment with 10 μM #20 followed by BNP stimulation. Data points are means±SEM (n=7).

### Enhanced potency of ANP-mediated vasorelaxation

Vasodilation contributes to the hypotensive properties of GC-A activation and can be modelled *ex vivo* in isolated rat aortic rings. To investigate the modulating effects of compounds #2 and #20, aortic rings that had been pre-contracted with U46619 were stimulated with increasing concentrations of these compounds (Fig. 6a) or with increasing concentrations of ANP in the presence or absence of 10 μM compound #2 or compound #20 (Fig. 6b). Compounds did not modulate vasorelaxation alone but reduced the EC_50_ of ANP from 4.7±0.4 nM to 2.2±0.2 nM (#2) and to 1.6±0.3 nM (#20).

**Fig. 6.**
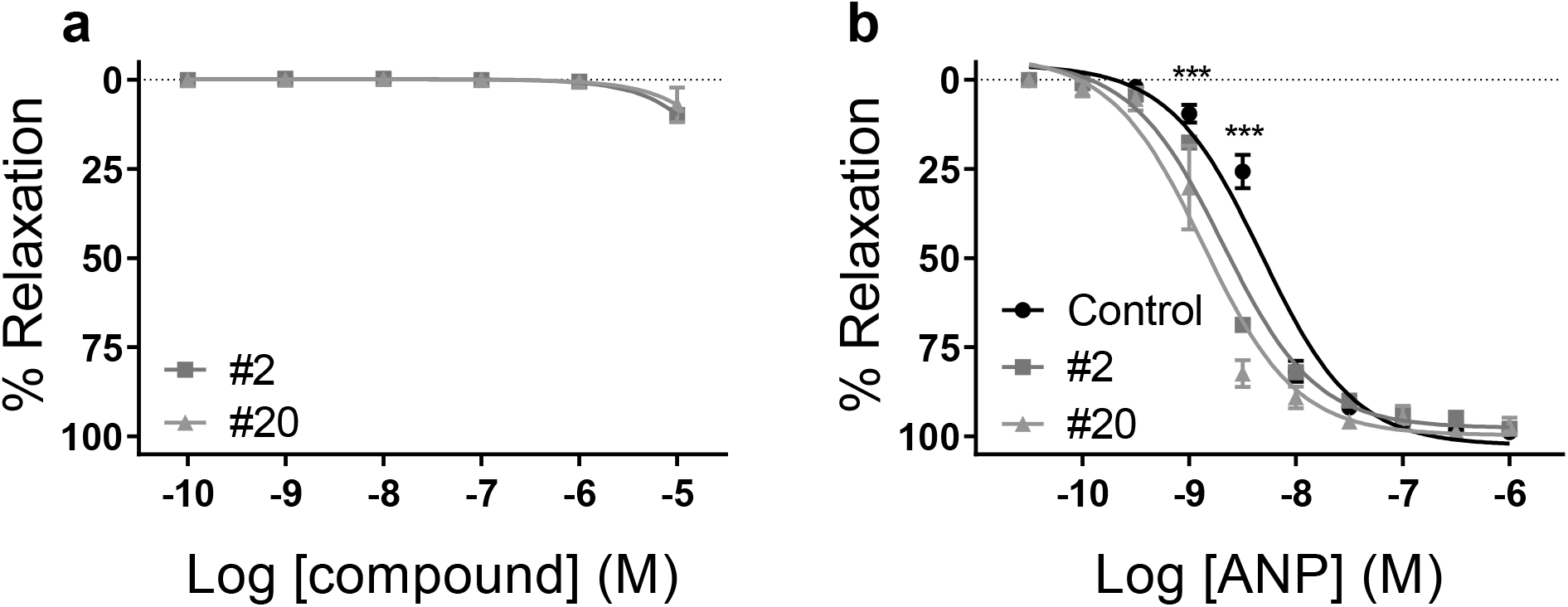
Compounds modulated the vasorelaxant activity of ANP but had no effect alone. Concentration-response of compounds #2 and #20 (**a**) and concentration-response of ANP in the absence or presence of 10 μM compound #2 or #20 (**b**) in isolated rat aorta pre-contracted with U46619 and in the presence of L-NAME. The relaxation is expressed as the means±SEM as a percentage of the U46619-induced tone. Data points are means±SEM (n=3-7). ***P<0.0001, two-way ANOVA (Bonferroni-corrected).

### Compound #20 increased the overall binding in whole cells

Binding assays that involved membranes from GC-A-expressing QBIHEK293A cells and ^125^I-ANP were first performed to investigate effects on binding. Both ANP and BNP were able to displace ^125^I-ANP from the receptor. However, there was no change in their binding or affinity in the presence of compound #2 or compound #20 (Fig. 7a,b). The compounds did not displace ^125^I-ANP or affect binding of ^125^I-ANP in concentrations up to 100 μM (Fig. 7c). However, when whole cells were used, compound #20 increased the overall binding of ANP by 60%±8% and slightly increased the IC_50_ from 2.3±0.3 nM to 3.0±0.4 nM (Fig. 7d). Increasing concentrations of compound #20 increased the binding of ^125^I-ANP by 71%±5% with an EC_50_ of 1.6±0.4 nM (Fig. 7e). In saturation binding assays, compound #20 did not change the affinity of ^125^I-ANP, but increased the maximal binding 2.4±0.1-fold (Fig. 7f).

**Fig. 7.**
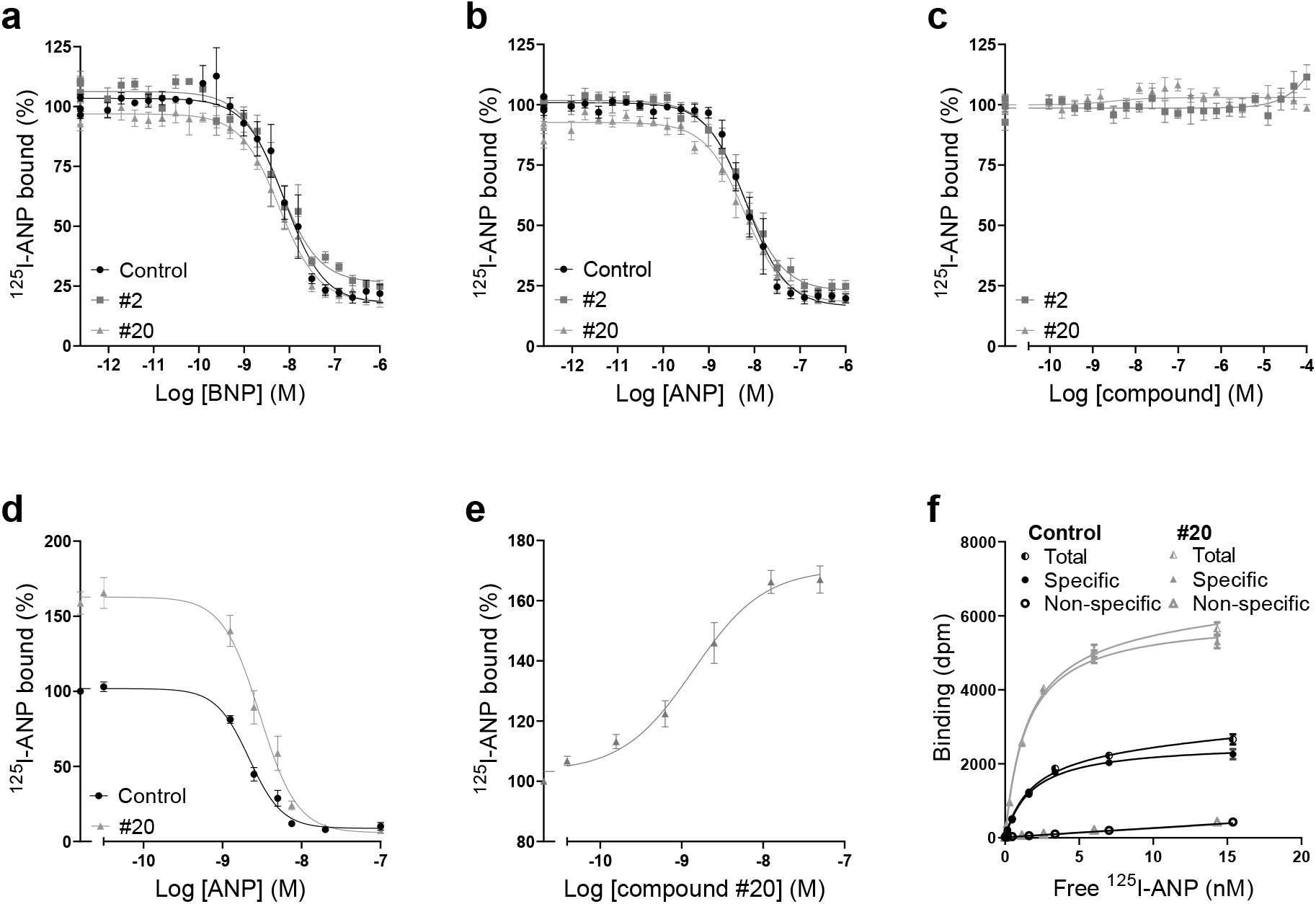
Compounds modulate binding, but not affinity, only in whole cells. **a-c,** Competition binding curves using membranes from GC-A-expressing cells with increasing concentrations of BNP (**a**) and ANP (**b**) with 50 pM ^125^I-ANP in the absence or presence of 10 μM compound #2 or #20 or with increasing concentrations of compounds #2 and #20 (**c**) (n=4-7). **d-f**, Binding curves using whole cells. Competition binding curves with increasing concentrations of ANP in the absence or presence of 10 μM compound #20 (**d**) and increasing concentrations of compound #20 (**e**). Data points are means±SEM (n=3-4). **f**, Saturation binding analysis of ^125^I-ANP in the presence (non-specific binding) or absence of 1 μM ANP (total binding), and in the absence or presence of 10 μM compound #20. Specific binding was determined by subtracting non-specific binding from total. Data points are means±SEM of triplicates from one representative assay of four assays performed in total.

### Compounds did not interfere with the allosteric binding sites for ATP on GC-A

Researchers have suggested that allosteric binding sites for ATP are present in the KHD and in the guanylyl cyclase domain. ATP binding to the KHD has been shown to reduce the affinity of NPs for GC-A^27^. The diuretic drug amiloride has been shown to antagonizes this effect of ATP by binding to the same site and increasing the affinity of ANP^27,28^. We wanted to explore whether compound #2 or #20 could antagonize the effect of ATP on the binding of ANP. In a competition binding assay with increasing concentrations of BNP, the presence of ATP reduced the binding of ^125^I-ANP, but co-incubation with the compounds had no additional effects on binding (Fig. 8a).

**Fig. 8.**
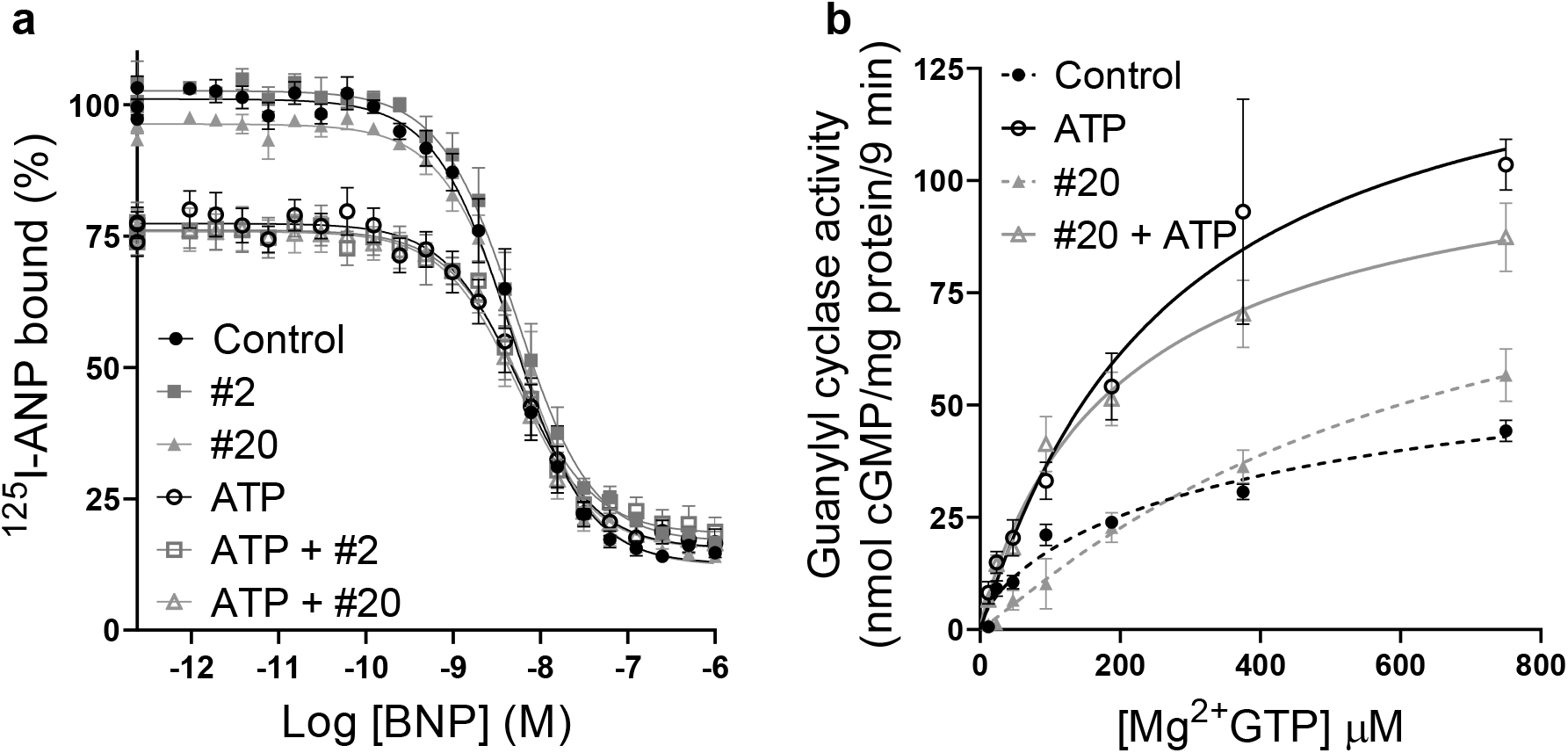
Compounds did not modulate the effects of ATP. **a,** Competition binding curves with increasing concentrations of cold BNP and 50 pM ^125^I-ANP in membranes from GC-A-expressing cells with or without 1 mM ATP or compound #2 or #20 (n=4). **b**, Substrate-velocity assay using membranes from HEK293T cells that expressed GC-A. The curves represent GC activity in the presence of 1 μM ANP with or without 1 mM ATP and/or compound #20 and the indicated Mg^2+^ GTP concentrations (n=3). Data points are means±SEM.

Previous studies have determined that ATP binds the pseudosymmetrical allosteric binding site in the catalytic domain of GC-A, and that this binding decreases the K_m_ of GTP from >1 mM to physiological concentrations of approximately 100 μM^29,46^. Importantly, in the absence of ATP, GTP binds to the allosteric site, which results in positive cooperativity. Here, we investigated the possibility that compound #20 activated GC-A by binding to the ATP allosteric site in the GC domain. However, in the absence of ATP, the presence of compound #20 did not change the K_m_ or cooperativity of the enzyme, which was inconsistent with the proposal that the compound bound to and activated this allosteric site (Fig. 8b).

### The effect of compound #20 is independent of phosphorylation of GC-A

Phosphorylation of multiple residues in the juxtamembrane and kinase homology domain is required for activation of GC-A. In human GC-A, there are seven phosphorylation sites^47^ and substitution of these sites with glutamate in GC-A^7E^ to mimic the negative charge of phosphate also mimics the phosphorylated and active form of GC-A^48^. By using GC-A^7E^, we investigated whether changes in phosphorylation contributed to the ability of compounds #20 and #2 to increase the activity of GC-A (Fig. 9). The presence of either compound increased the level of ANP- or BNP-mediated cGMP further in GC-A^7E^ by 151%±23% (#2) and 183±26% (#20) for ANP and 170%±25% (#2) and 221%±33% (#20) for BNP. The compounds also decreased the EC_50_ for both ANP (1.8±0.2-fold (#2) and 1.6±0.2-fold (#20)) and BNP (2.9±0.6-fold (#2) and 3.9±1.1-fold (#20)).

**Fig. 9.**
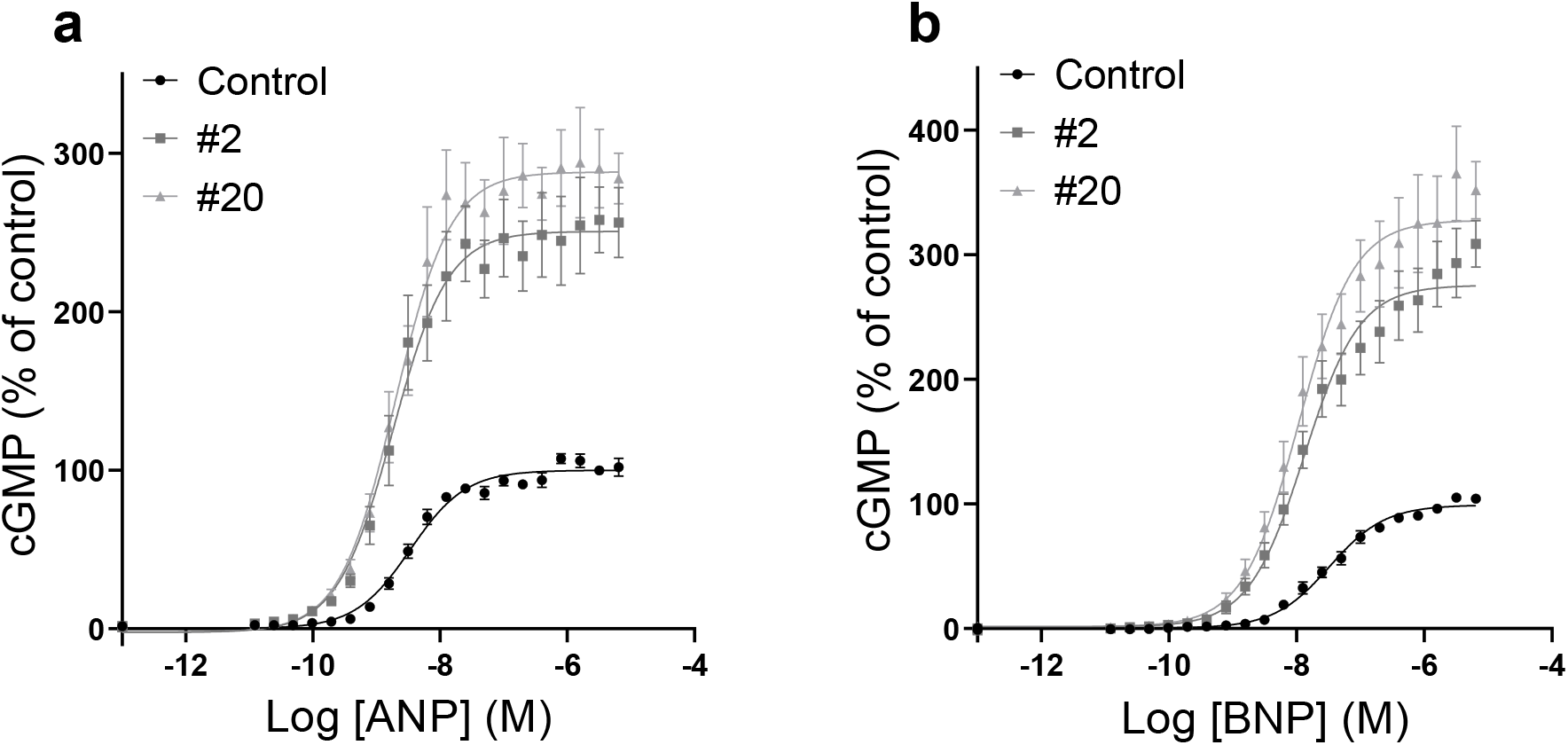
Phosphorylation state of GC-A does not prevent the activity of compounds. Concentration-response curves for stimulation by ANP (**a**) or BNP (**b**) of cGMP production by the phosphomimetic mutant GC-A^7E^ in the presence or absence of 10 μM compound #2 or #20. Data points are means±SEM (n=5-7).

### No modulation of chaperone activity towards GC-A

GC-A can be modulated indirectly by proteins that facilitate the correct folding of the enzyme. It has been shown that heat shock protein 90 (HSP90) is a chaperone that interacts with GC-A, and that inhibition of HSP90 with geldanamycin reduces levels of ANP-mediated cGMP production^49^. To investigate whether our compounds were involved in HSP90 folding and trafficking of GC-A, we inhibited HSP90 and measured levels of ANP- and BNP-mediated cGMP production with and without the presence of compound #20 (Fig. 10). As in the previous study, inhibition of HSP90 reduced ANP- and BNP-mediated cGMP production by 45%±11% and 41%±12%, respectively. We did not observe a change in the potency of ANP or BNP. When we inhibited HSP90, compound #20 increased the NP-mediated cGMP production and reduced the EC_50_ of ANP and BNP similar to what we observed in the absence of HSP90 inhibition. The presence of compound #20 increased the cGMP production by 7%±6% (ANP) and 11%±5% (BNP), and reduced the EC_50_ by 1.7±0.5-fold and 3.6±1.6-fold for ANP and BNP, respectively.

**Fig. 10.**
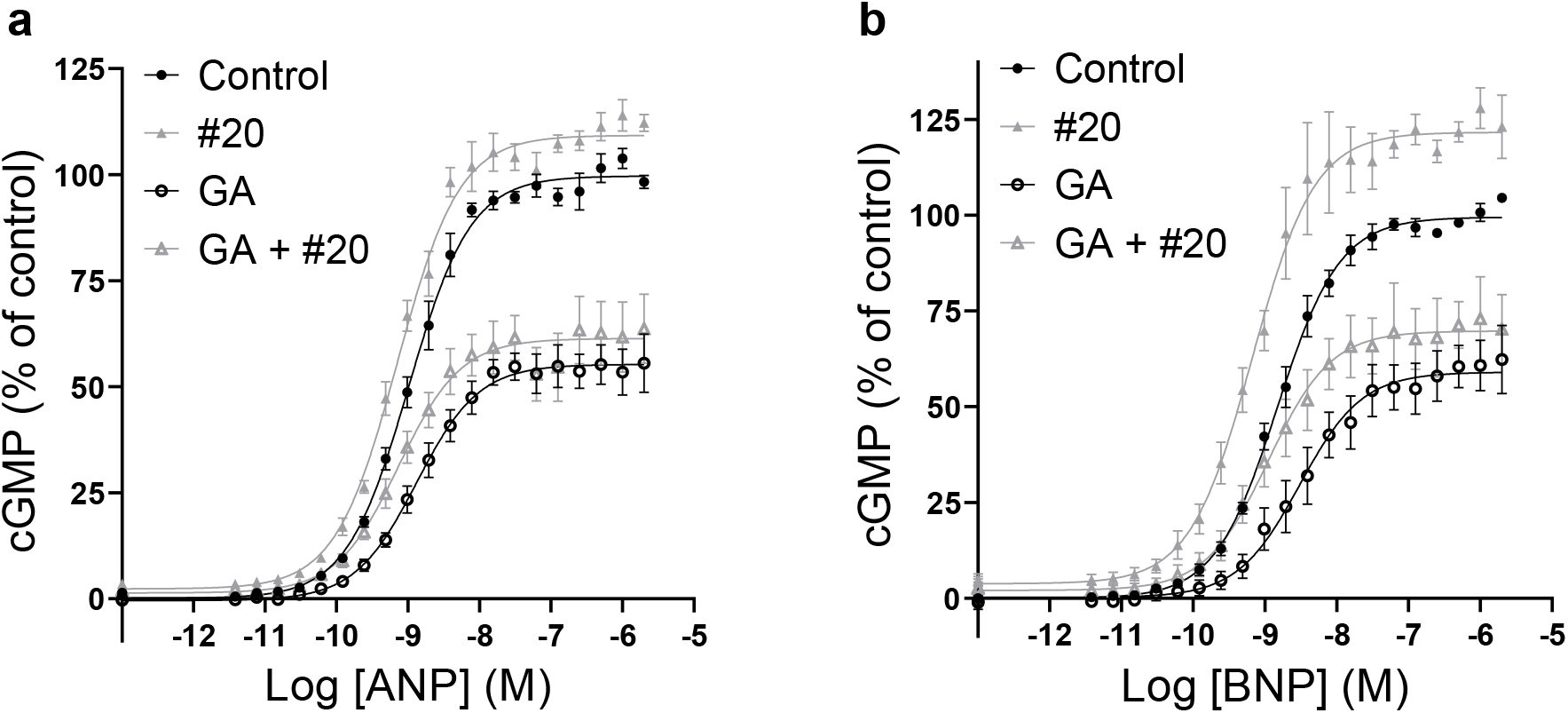
Compounds work independently of HSP90 activity. ANP-stimulated (**a**) and BNP-stimulated (**b**) cGMP production in cells that expressed GC-A in the presence or absence of 10 μM compound #20 and the chaperone HSP90 inhibitor geldanamycin (GA; 10 μM). Data points are means±SEM (n=4-8).

### The effect of compound #20 follows the intracellular domain of GC-A

Human GC-A and GC-B are 57% identical in the extracellular domain and 78% identical in the intracellular domain. The dissimilarity in the extracellular domain is thought to explain their different affinities for NPs, while their intracellular domains are highly conserved, especially the guanylyl cyclase domain, which is 92% identical at the amino acid level. Compound #20 seemed to be highly selective towards GC-A, with no effects towards GC-B or NPR-C. By making chimeric GC-A/GC-B in which domains, regions, and amino acids had been swapped between the two receptors, we could investigate which part of GC-A affected the activity of compound #20. The extracellular domain of GC-A comprises approximately half of the receptor (amino acids 33-473), followed by a short 21-amino acid transmembrane domain (amino acids 474-494) and an intracellular domain of 567 amino acids from 496-1061. When the intracellular domain of GC-A was replaced with the intracellular domain of GC-B (GC-A^1-494^/B^479-1047^), compound #20 became inactive (Fig. 11a). Conversely, for the analogous GC-B^1-478^/A^495-1061^, the presence of compound #20 led to a change in the EC_50_ of CNP towards lower concentrations, but it did not increase the maximum CNP-mediated cGMP production.

**Fig. 11.**
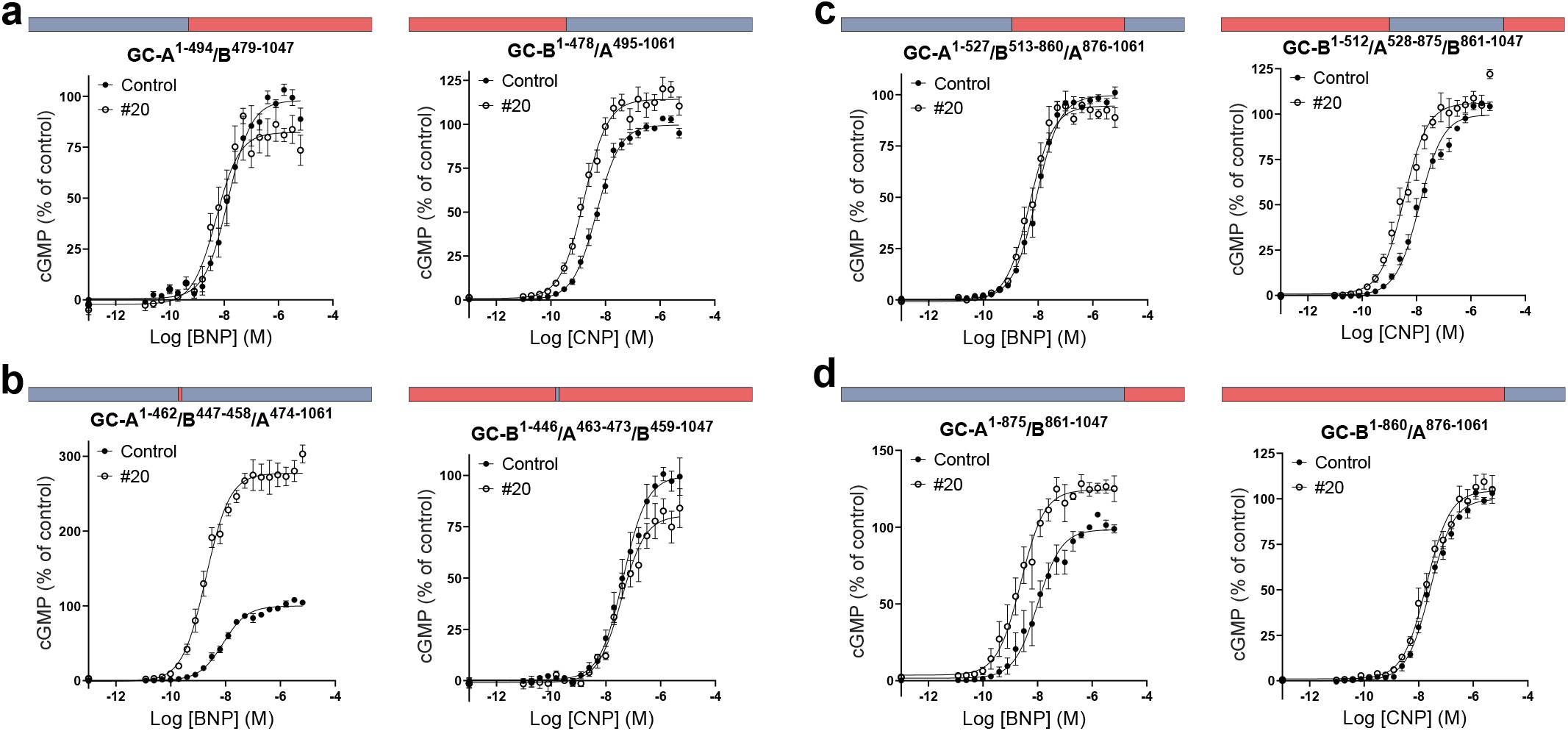
The activity of compound #20 followed the intracellular domain. Chimeric GC-A (blue)/GC-B (red) pairs stimulated with either increasing concentrations of BNP (GC-A extracellular domain) or CNP (GC-B extracellular domain) in the presence of 0.1% DMSO in control or 10 μM compound #20. The graphs show the activity of compound #20 after swapping the intracellular domain (**a**), the juxtamembrane domain (**b),** the kinase homology domain and the coiled-coil domain (**c**) and the guanylyl cyclase domain (**d**) between GC-A and GC-B. Data points are means±SEM (n=4-8).

We were also interested in investigating a previously described allosteric binding site for the unselective antagonist HS-142-1 in the juxtamembrane domain in GC-A and GC-B^26^. When only the juxtamembrane domain was swapped between the two receptors, the selectivity of compound #20 towards GC-A^1-462^/B^447-458^/A^474-1061^ was maintained (Fig. 11b). Further, when the KHD and the CCD were swapped between the two receptors, the activity of compound #20 moved to GC-B^1-512^/A^528-875^/B^861-1047^ and the activity towards GC-A^1-527^/B^513-860^/A^876-1061^ was prevented (Fig. 11c). Swapping only the KHDs or the CCDs between the two receptors yielded non-functional GC-B^1-512^/A^528-805^/B^787-1047^ and GC-A^1-805^/B^787-860^/A^876-1061^, respectively. The activity of compound #20 was conserved between GC-A^1-875^/B^861-1047^ and GC-B^1-860^/A^876-1061^ where only the guanylyl cyclase domains were swapped between the two receptors (Fig. 11d).

### The effect of compound #20 depends on the amino acid residues at position 640 in GC-A

Since the activity of compound #20 seemed to be determined by the KHD and/or CCD, we constructed and tested several chimeric receptors that were focused on this region from amino acids 528 to 875 in GC-A (results summarized in Fig. 12, and all graphs are shown in Supplementary Fig. S2). It became apparent that the activity of compound #20 followed the region from amino acids 621 to 729 in GC-A, since most of the activity of compound #20 was lost for GC-A^1-620^/B^605-714^/A^730-1061^ but gained in GC-B^1-604^/A^621-729^/B^715-1047^ (Fig. 13a). Further, the activity was lost in GC-A^1-621^/B^605-647^/A^664-1061^ and present in GC-B^1-604^/A^621-663^/B^648-1047^ (Fig. 13b), whereas no change in activity was observed for GC-A^1-663^/B^648-686^/A^701-1061^ and the corresponding GC-B^1-647^/A^664-700^/B^686-1047^ (13c) or for GC-A^1-700^/B^686-715^/A^731-1061^ and the corresponding GC-B^1-685^/A^701-730^/B^716-1047^ (13d). The amino acid sequence from 621 to 663 in GC-A is highly conserved in GC-B and only nine amino acids differ between the two receptors. We constructed chimeric receptors that involved the switching of one or two amino acids between the two receptors (all graphs shown in Supplementary Fig. S3) and saw that the activity of compound #20 only followed amino acid Thr640 in GC-A. Compound #20 became completely inactive towards GC-A^T640I^ but it gained activity towards GC-B^I624T^ (Fig. 14a). By substituting Ile624 with Thr in GC-B, compound #20 increased the maximum CNP-mediated cGMP production by 35%±6% and decreased the EC_50_ for CNP 4.4±0.7-fold. However, the activity of compound #20 was not exclusively dependent on the presence of Thr. Compound #20 was also active when GC-A^T640^ or GC-B^I624^ were substituted with Ala and Ser (Fig. 14c,e), but not when they were substituted with Tyr and Leu (Fig. 14b,f). Compound #20 was not active in GC-A^T640V^, but was active in GC-B^I624V^ (14d). Substitution of Thr640 (GC-A) or Ile624 (GC-B) with glutamic acid or aspartic acid produced non-functional receptors.

**Fig. 12.**
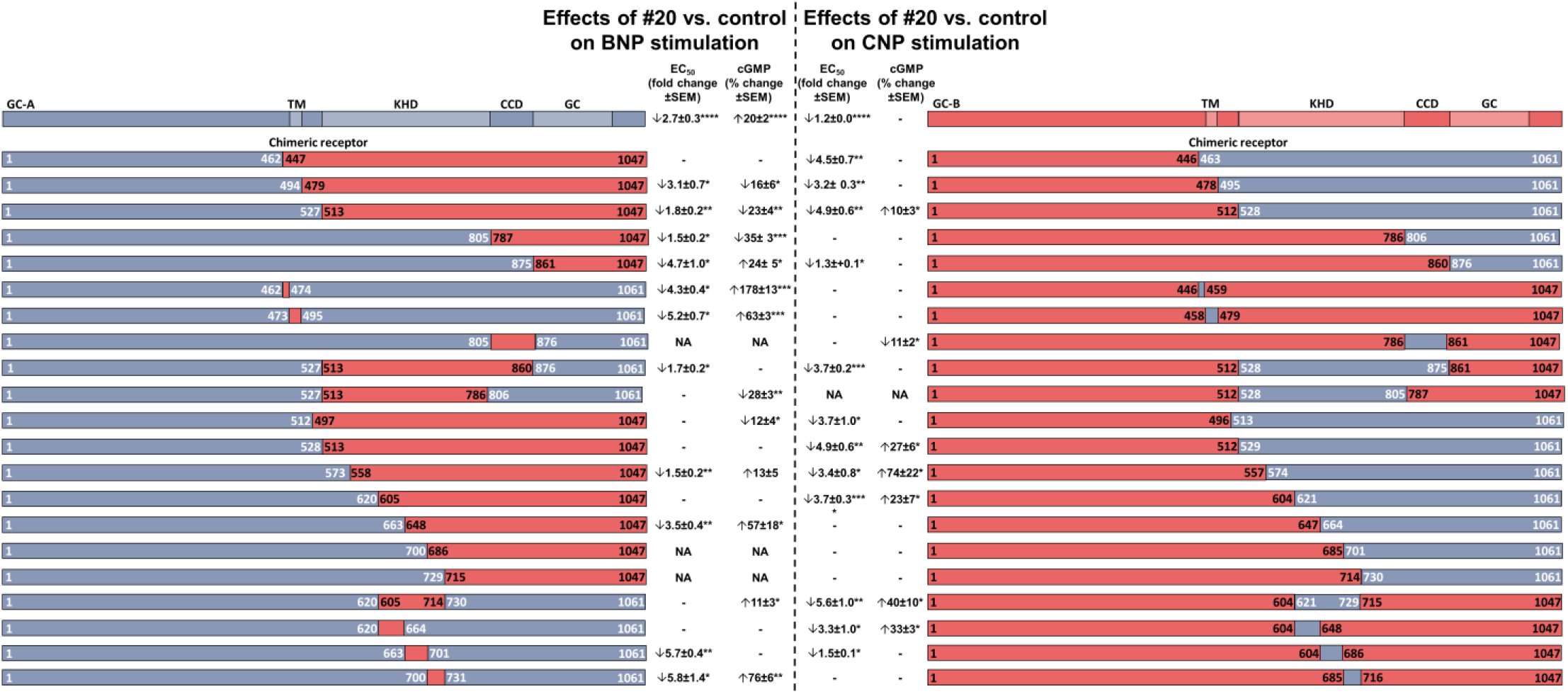
Illustrative summary of the results from all tested chimeric GC-A/B receptors. The effect of compound #20 versus control on BNP or CNP stimulation towards chimeric GC-A/B. The effects of compound #20 are quantified as fold change in EC_50_±SEM and percentage change±SEM in the maximal NP-mediated cGMP production. Compound #20 increased (↑), decreased (↓) or had no effect (-) on the EC_50_ or cGMP production during testing of the illustrated chimeric GC-A/B. Some chimeric receptors were not active (NA) in response to NP stimulation. Effects were analyzed as difference between control and compound #20 and validated using t-test. *p≤0.05, **p≤0.01, ***p≤0.001. The individual graphs are shown in Supplementary Fig. S2. TM, transmembrane domain; KHD, kinase homology domain; CCD, dimerization domain; GC, guanylyl cyclase domain.

**Fig. 13.**
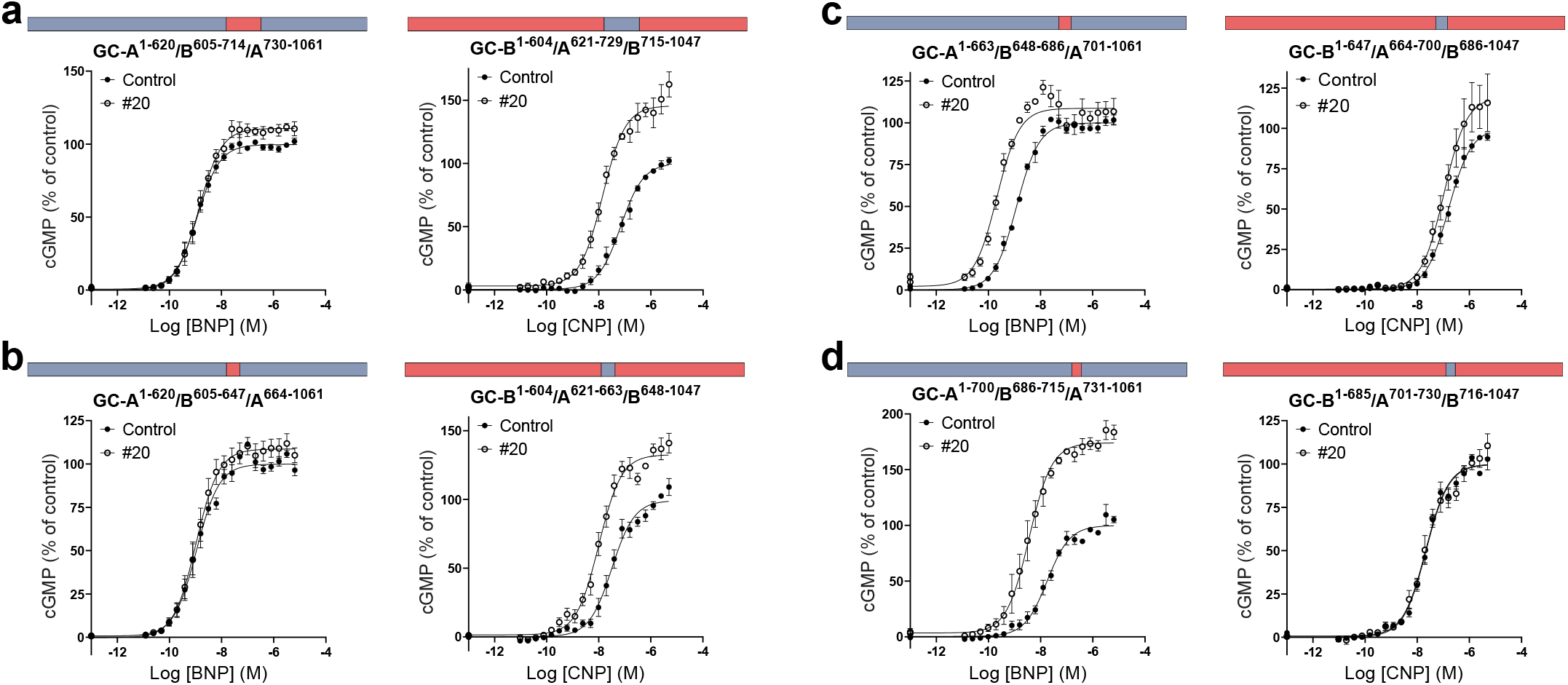
The activity of compound #20 depended on the presence of a small region in the kinase homology domain. Chimeric receptors with focus on the kinase homology domain and the dimerization domain showed that compound #20 was active only when the region of amino acid residues 621-661 was present. The activity moved to GC-B when the corresponding amino acid residues were replaced with those of GC-A^621-729^ (**a**) or GC-A^621-663^ (**b**), but not GC-A^664-700^ (**c**) or GC-A^701-730^ (**d**). Data points are means±SEM (n=4-8).

**Fig. 14.**
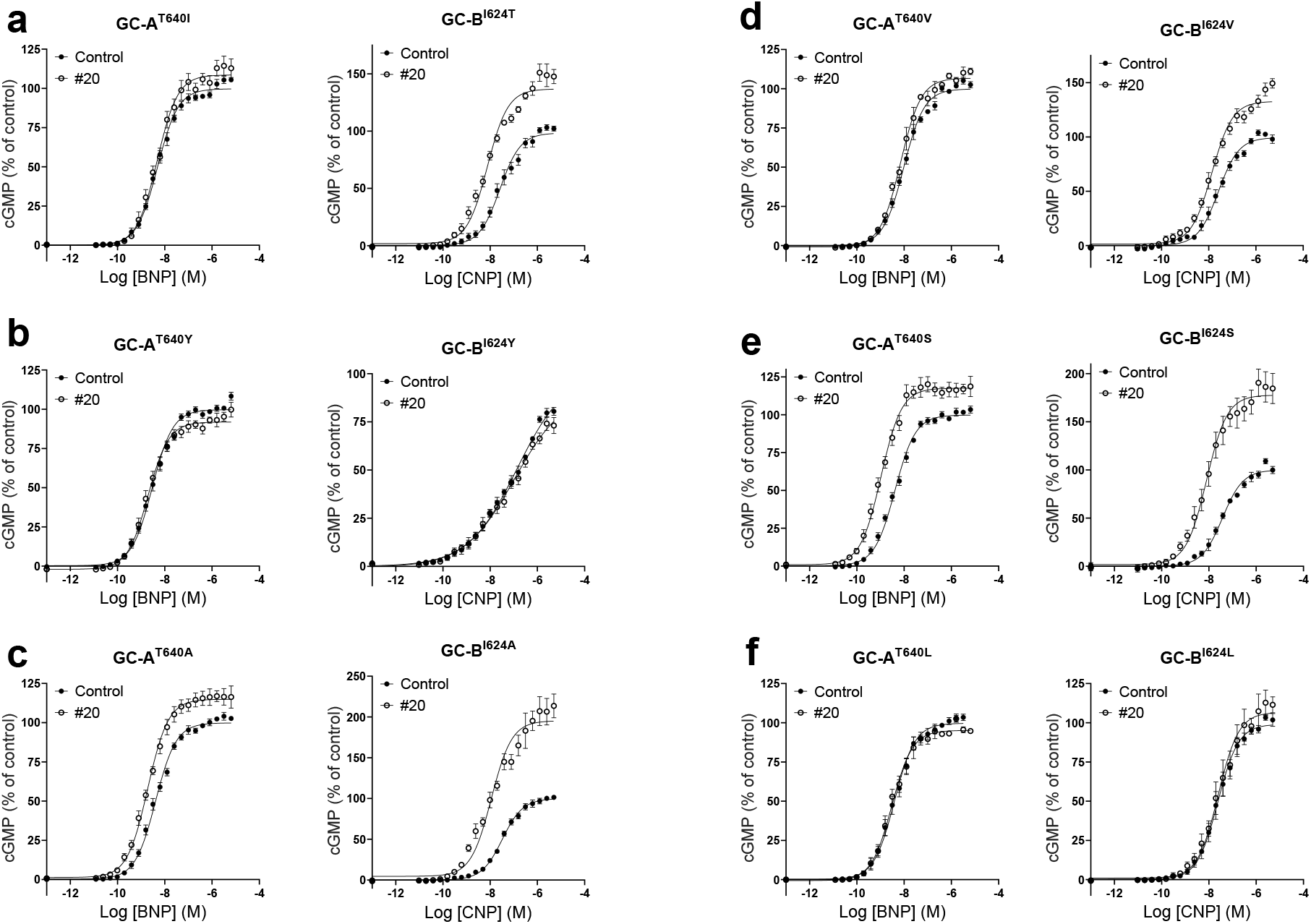
Only threonine 640 mediates the effect of compound #20. **a,** Swapping of nine non-conserved amino acids in the region 621-663 between GC-A and GC-B revealed that the activity of compound #20 was only dependent on the presence of threonine 640 in GC-A. **b-f**, Substitution of GC-A^T640^ and the corresponding GC-B^1624^ with the amino acids tyrosine, alanine, valine, serine and leucine. Data points are means±SEM (n=4-8).

The structure of the intracellular domains of GC-A or GC-B have not yet been solved, but homology models of their KHDs have been published^32^. These models were based on conserved structure similarities between KHDs in GC-A and GC-B and protein kinases. In addition, the structures of the full-length receptors have been predicted using deep-learning system AlphaFold^50^. In both models, GC-A^T640^ and GC-B^I624^ are buried in an alpha helical region and are not readily accessible from the surface (Fig. 15).

**Fig. 15.**
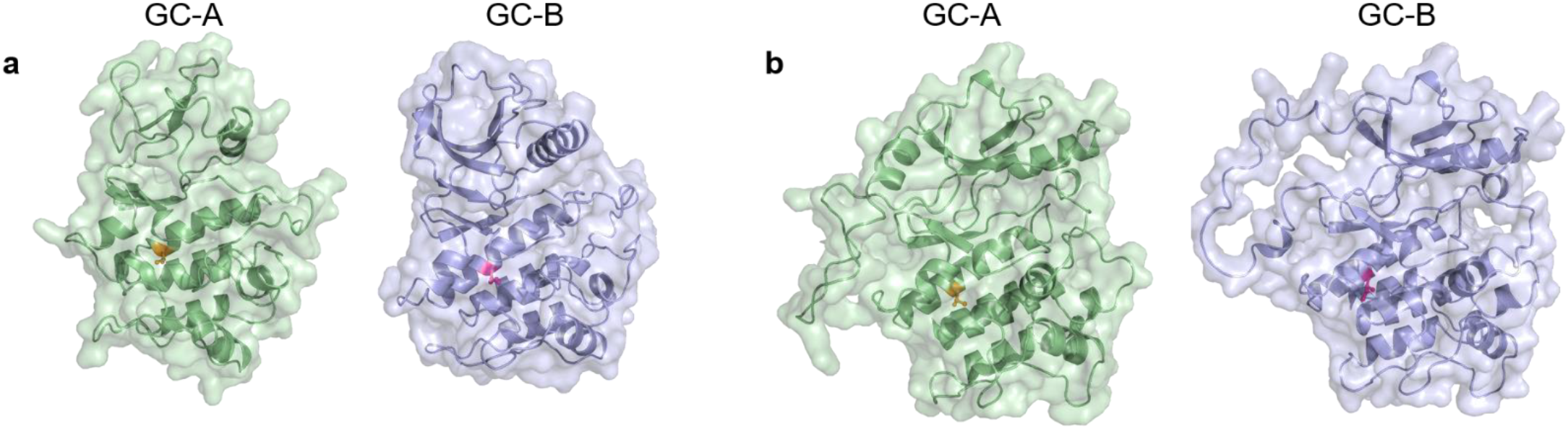
GC-A^T640^ and GC-B^1624^ are buried in an alpha helical region. Homology models (**a**) and predicted models (**b**) of GC-A and GC-B in which GC-A^T640^ is in yellow and GC-B^1624^ is in pink.

## DISCUSSION

Here we show that small molecular compounds can increase the BNP- or ANP-mediated cGMP production and thereby enhance the efficacy of these peptides towards GC-A. We further identified a key amino acid in GC-A, the presence of which is necessary for this effect.

### Small molecular allosteric enhancers of GC-A

GC-A is an attractive target for the treatment of cardiovascular diseases because its activation leads to vasodilation, and has an overall cardiorenal protective effect. The current strategy is to use BNP, ANP and designer NPs, but their inherently short half-life and poor bioavailability limit their potential uses. Another strategy is to increase their concentrations by inhibiting neprilysin and hence preventing their catabolism. However, neprilysin is not selective towards NPs. It also catabolizes angiotensin II and other hormones that counteract the effects of NPs.

In contrast with previous strategies, small molecular modulators uniquely enhance the effects of endogenous ANP and BNP towards GC-A. Modulation of allosteric binding sites can have several advantages besides their lack of competition with NPs towards their binding site. Allosteric binding sites are thought to be less conserved than orthosteric sites, so there is opportunity to use more selective compounds^51^. We have shown that the activity and selectivity of compound #20 depend on the presence of one non-conserved residue between GC-A and GC-B. In addition, allosteric modulators can be safer than orthosteric agonists, since their effects are limited to both the presence and the concentration of the endogenous agonist. Too much GC-A activation can lead to hypotension and was shown to be a safety concern when using BNP in treating heart failure^1^.

### Novel allosteric binding site on GC-A

In our search for the allosteric binding site of our compounds, we started to investigate known allosteric binding sites in the extracellular and intracellular domains of GC-A. Our GC-A/B chimeric constructs excluded the possibility that the sought site was one of two known extracellular allosteric binding sites on GC-A (Fig. 11a,b). The first of these was a site for a chloride atom in the extracellular domain that regulates binding of ANP to GC-A and has been suggested to be an allosteric modulator in the kidneys^24,25^. The second was the putative allosteric binding site for the unselective GC-A antagonist HS-142-1 in the juxtamembrane domain^26^. In the intracellular domain, the chimeric receptors also excluded the possibility that our compounds were bound to the allosteric binding sites that have been suggested for ATP in the KHD and GC domains^27–29,46^. Testing of more chimeric GC-A/B receptors narrowed the search for the putative binding site to a region in the KHD of GC-A and finally pinpointed a single amino acid. However, the activity of compound #20 at the chimeric receptors was not always similar to that at GC-A. For some chimeric receptors, the presence of compound #20 affected only the EC_50_ or the maximum cGMP level but not both. This could reflect differences in receptor expression levels and receptor reserve in the transfected cells^52^. In a few chimeric pairs, compound #20 was active at both, but more at the chimeric receptors containing Thr640. The activity of compound #20 followed that of GC-A^T640^ in all chimeric pairs, except for GC-A^1-805^/B^787-1047^ and the corresponding GC-B^1-786^/A^806-1061^, in which it reduced the BNP-mediated cGMP production by GC-A^1-805^/B^787-1047^. These deviating results might imply that the activity of compound #20 on these chimeric GC-A/B receptors could be affected by intramolecular interactions between homologous regions of GC-A and GC-B. However, no amino acid substitution other than that of GC-A^T640^ prevented the activity, and activity in GC-B was restored by substitution of the analogous GC-B^I624T^.

Our chimeric GC-A/B receptors and point mutations do not provide definite evidence of a binding site. Mutations in one part of GC-A or GC-B can affect the function of the receptor elsewhere through intramolecular interactions. Also, the binding site may be located at a different site on the receptor, but the effect be mediated through GC-A^T640^. From the existing models of GC-A and GC-B, we see that both GC-A^T640^ and GC-B^I624^ are buried in an alpha helix (Fig. 15) and may not be readily accessible for a small molecule to interact with. We also lack evidence of a direct interaction between compound #20 and GC-A. This could be investigated further through resolution of the structure of GC-A with compound #20.

### Unknown mechanism of action

Allosteric enhancers are compounds that enhance the affinity and/or efficacy of the orthosteric agonist while having no effect on their own^53^. In our concentration-response curves, we did not detect any effects of the compounds alone on cGMP production (Figs. 2a and 4a), vasorelaxation (Fig. 6a) or on basal GC-A activity (Fig. 5). Our binding experiments showed that the compounds did not affect the affinities of ANP or BNP in either whole cells or membranes, but increased the overall binding in whole cells (Fig. 7). Therefore, we explored the known mechanisms of action of allosteric modulators of GC-A, which modulate cGMP production and the efficacy of NPs. The presence of ATP increases the efficacy of the catalytic guanylyl cyclase domain by decreasing K_m_ and increasing V_max_^29,30^, but it has also been shown to reduce the overall binding of ANP to GC-A^27,28^. However, none of our compounds modulated the effects of ATP on binding and they did not change the efficacy of the enzyme in our GC assays with or without ATP present (Fig. 8a,b). We also investigated known methods of indirect modulation of GC-A and found that our compounds did not affect the effect of HSP90 or increase cGMP production by inhibiting PDEs. These findings indicate that our compounds allosterically enhance the efficacy of NPs through a currently unknown mechanism of action.

To identify small molecules that activated GC-A, we performed a high throughput screening. Through this process, only one compound was identified as a hit. Small molecular GC-A agonists have been described previously^6–8^, but to our knowledge, identification of allosteric enhancers of GC-A has not been described. Our low hit rate and the fact that only a few small compounds are known to activate GC-A could be due to the structure and large interface of the orthosteric binding site. The use of small molecules to mimic NP activation of GC-A remains a challenge in the utilization of GC-A as a potential drug target. Here, we provide the structure and activity of two allosteric enhancers of GC-A and suggest a novel allosteric binding site. Both compounds could serve as tool compounds for further development and proof-of-concept of allosteric enhancement of GC-A.

## ACKNOWLEDGEMENTS

This work was supported by the South-Eastern Norway Regional Health Authority (grants 2019051 and 30245), the Research Council of Norway (grants 205167 and 303490), the Anders Jahre Foundation for the Promotion of Science, the Family Blix Foundation, grants from the University of Oslo, grant from the Norwegian Health Association (grant 460310) and a grant from the Norwegian Centennial Chair programme. The authors would like to acknowledge the help of Kristin Nordskogen Smeby and Iwona Gutowska-Schiander in the conduct of some of the experiments.

## AUTHOR CONTRIBUTIONSs

All authors designed the research. CPT, JR, DD, AJH, LRM and HA performed research and analyzed data. HA wrote the manuscript with input from all authors. All authors approved the final version.

## REFERENCES

1 O’Connor, C. M. et al. Effect of Nesiritide in Patients with Acute Decompensated Heart Failure. New England Journal of Medicine 365, 32–43, doi:10.1056/NEJMoa1100171 (2011).

2 Packer, M. et al. Effect of Ularitide on Cardiovascular Mortality in Acute Heart Failure. New England Journal of Medicine 376, 1956–1964, doi:10.1056/NEJMoa1601895 (2017).

3 Kawakami, R. et al. A Human Study to Evaluate Safety, Tolerability, and Cyclic GMP Activating Properties of Cenderitide in Subjects With Stable Chronic Heart Failure. Clin Pharmacol Ther 104, 546–552, doi: 10.1002/cpt.974 (2018).

4 Cataliotti, A., Costello-Boerrigter, L. C., Chen, H. H., Textor, S. C. & Burnett, J. C., Jr. Sustained blood pressure-lowering actions of subcutaneous B-type natriuretic peptide (nesiritide) in a patient with uncontrolled hypertension. Mayo Clinic proceedings 87, 413–415, doi:10.1016/j.mayocp.2012.02.003 (2012).

5 Chen, Y. et al. Long-term blood pressure lowering and cGMP-activating actions of the novel ANP analog MANP. American journal of physiology. Regulatory, integrative and comparative physiology 318, R669–r676, doi: 10.1152/ajpregu.00354.2019 (2020).

6 Iwaki, T. et al. Discovery and dimeric approach of novel Natriuretic Peptide Receptor A (NPR-A) agonists. Bioorganic & medicinal chemistry 25, 1762–1769, doi:https://doi.org/10.1016/j.bmc.2017.01.026 (2017).

7 Iwaki, T. et al. Discovery and SAR of a novel series of Natriuretic Peptide Receptor-A (NPR-A) agonists. Bioorganic & medicinal chemistry letters 27, 4904–4907, doi:https://doi.org/10.1016/j.bmcl.2017.09.028 (2017).

8 Iwaki, T. et al. Discovery and in vivo effects of novel human natriuretic peptide receptor A (NPR-A) agonists with improved activity for rat NPR-A. Bioorganic & medicinal chemistry 25, 6680–6694, doi:https://doi.org/10.1016/j.bmc.2017.11.006 (2017).

9 Potter LR, A.-H. S., Dickey DM. Natriuretic Peptides, Their Receptors, and Cyclic Guanosine Monophosphate-Dependent Signaling Functions. Endocrine Reviews 27, 47–72, doi:doi:10.1210/er.2005-0014 (2006).

10 Holtwick, R. et al. Smooth muscle-selective deletion of guanylyl cyclase-A prevents the acute but not chronic effects of ANP on blood pressure. Proc Natl Acad Sci U S A 99, 7142–7147, doi:10.1073/pnas.102650499 (2002).

11 Lopez, M. J., Garbers, D. L. & Kuhn, M. The guanylyl cyclase-deficient mouse defines differential pathways of natriuretic peptide signaling. The Journal of biological chemistry 272, 23064–23068, doi:10.1074/jbc.272.37.23064 (1997).

12 Lopez, M. J. et al. Salt-resistant hypertension in mice lacking the guanylyl cyclase-A receptor for atrial natriuretic peptide. Nature 378, 65–68, doi:10.1038/378065a0 (1995).

13 Oliver, P. M. et al. Hypertension, cardiac hypertrophy, and sudden death in mice lacking natriuretic peptide receptor A. Proc Natl Acad Sci U S A 94, 14730–14735 (1997).

14 John, S. W. et al. Genetic decreases in atrial natriuretic peptide and salt-sensitive hypertension. Science 267, 679–681 (1995).

15 John, S. W. et al. Blood pressure and fluid-electrolyte balance in mice with reduced or absent ANP. The American journal of physiology 271, R109–114 (1996).

16 Holditch, S. J. et al. B-Type Natriuretic Peptide Deletion Leads to Progressive Hypertension, Associated Organ Damage, and Reduced Survival: Novel Model for Human Hypertension. Hypertension 66, 199–210, doi:10.1161/hypertensionaha.115.05610 (2015).

17 Vandenwijngaert, S. et al. Blood Pressure-Associated Genetic Variants in the Natriuretic Peptide Receptor 1 Gene Modulate Guanylate Cyclase Activity. Circ Genom Precis Med 12, e002472, doi:10.1161/circgen.119.002472 (2019).

18 Nakayama, T. et al. Functional deletion mutation of the 5’-flanking region of type A human natriuretic peptide receptor gene and its association with essential hypertension and left ventricular hypertrophy in the Japanese. Circ Res 86, 841–845 (2000).

19 Belluardo, P. et al. Lack of activation of molecular forms of the BNP system in human grade 1 hypertension and relationship to cardiac hypertrophy. American journal of physiology. Heart and circulatory physiology 291, H1529–1535, doi:10.1152/ajpheart.00107.2006 (2006).

20 Newton-Cheh, C. et al. Association of common variants in NPPA and NPPB with circulating natriuretic peptides and blood pressure. Nature genetics 41, 348–353, doi:10.1038/ng.328 (2009).

21 He, X.-l., Chow, D.-c., Martick, M. M. & Christopher Garcia, K. Allosteric Activation of a Spring-Loaded Natriuretic Peptide Receptor Dimer by Hormone. Science 293, 1657–1662, doi:10.1126/science.1062246 (2001).

22 Ogawa, H., Qiu, Y., Ogata, C. M. & Misono, K. S. Crystal Structure of Hormone-bound Atrial Natriuretic Peptide Receptor Extracellular Domain: ROTATION MECHANISM FOR TRANSMEMBRANE SIGNAL TRANSDUCTION. Journal of Biological Chemistry 279, 28625–28631, doi:10.1074/jbc.M313222200 (2004).

23 Misono, K. S. et al. Structure, signaling mechanism and regulation of natriuretic peptide receptor-guanylate cyclase. The FEBS journal 278, 1818–1829, doi:10.1111/j.1742-4658.2011.08083.x (2011).

24 Misono, K. S. Atrial Natriuretic Factor Binding to Its Receptor Is Dependent on Chloride Concentration: A Possible Feedback-Control Mechanism in Renal Salt Regulation. Circulation Research 86, 1135–1139, doi: 10.1161/01.res.86.11.1135 (2000).

25 Ogawa, H. et al. Reversibly bound chloride in the atrial natriuretic peptide receptor hormone-binding domain: possible allosteric regulation and a conserved structural motif for the chloride-binding site. Protein science : a publication of the Protein Society 19, 544–557, doi:10.1002/pro.332 (2010).

26 Poirier, H., Labrecque, J., Deschenes, J. & DeLean, A. Allotopic antagonism of the non-peptide atrial natriuretic peptide (ANP) antagonist HS-142-1 on natriuretic peptide receptor NPR-A. Biochem J 362, 231–237 (2002).

27 Jewett, J. R., Koller, K. J., Goeddel, D. V. & Lowe, D. G. Hormonal induction of low affinity receptor guanylyl cyclase. The EMBO journal 12, 769–777 (1993).

28 De Lean, A. Amiloride potentiates atrial natriuretic factor inhibitory action by increasing receptor binding in bovine adrenal zona glomerulosa. Life sciences 39, 1109–1116 (1986).

29 Robinson, J. W. & Potter, L. R. Guanylyl Cyclases A and B Are Asymmetric Dimers That Are Allosterically Activated by ATP Binding to the Catalytic Domain. Science signaling 5, ra65–ra65, doi:10.1126/scisignal.2003253 (2012).

30 Robinson, J. W. & Potter, L. R. ATP potentiates competitive inhibition of guanylyl cyclase A and B by the staurosporine analog, Go6976: reciprocal regulation of ATP and GTP binding. The Journal of biological chemistry 286, 33841–33844, doi: 10.1074/jbc.M111.273565 (2011).

31 Potter, L. R. & Hunter, T. Phosphorylation of the kinase homology domain is essential for activation of the A-type natriuretic peptide receptor. Mol Cell Biol 18, 2164–2172 (1998).

32 Edmund, A. B., Walseth, T. F., Levinson, N. M. & Potter, L. R. The pseudokinase domains of guanylyl cyclase-A and -B allosterically increase the affinity of their catalytic domains for substrate. Science signaling 12, doi:10.1126/scisignal.aau5378 (2019).

33 Heim, J.-M., Singh, S. & Gerzer, R. Effect of glycosylation on cloned ANF-sensitive guanylyl cyclase. Life sciences 59, PL61–PL68, doi:https://doi.org/10.1016/0024-3205(96)00306-2 (1996).

34 Tanaka, T. D., Sawano, M., Ramani, R., Friedman, M. & Kohsaka, S. Acute heart failure management in the USA and Japan: overview of practice patterns and review of evidence. ESC Heart Failure 5, 931–947, doi:https://doi.org/10.1002/ehf2.12305 (2018).

35 Potter, L. R. Natriuretic peptide metabolism, clearance and degradation. FEBS Journal 278, 1808–1817, doi:10.1111/j.1742-4658.2011.08082.x (2011).

36 Dickey, D. M. & Potter, L. R. Dendroaspis natriuretic peptide and the designer natriuretic peptide, CD-NP, are resistant to proteolytic inactivation. J Mol Cell Cardiol 51, 67–71, doi:10.1016/j.yjmcc.2011.03.013 (2011).

37 Chen, Y. et al. CRRL269: A Novel Designer and Renal Enhancing pGC-A Peptide Activator. American journal of physiology. Regulatory, integrative and comparative physiology, ajpregu.00286.02017, doi:10.1152/ajpregu.00286.2017 (2017).

38 McMurray, J. J. et al. Angiotensin-neprilysin inhibition versus enalapril in heart failure. The New England journal of medicine 371, 993–1004, doi:10.1056/NEJMoa1409077 (2014).

39 Bach, T. et al. Identification of small molecule NPR-B antagonists by high throughput screening — potential use in heart failure. Naunyn-Schmiedeberg’s Arch Pharmacol 387, 5–14, doi:10.1007/s00210-013-0940-6 (2014).

40 Dickey, D. M., Yoder, A. R. & Potter, L. R. A familial mutation renders atrial natriuretic Peptide resistant to proteolytic degradation. The Journal of biological chemistry 284, 19196–19202, doi:10.1074/jbc.M109.010777 (2009).

41 Marchmont, R. J. & Houslay, M. D. A peripheral and an intrinsic enzyme constitute the cyclic AMP phosphodiesterase activity of rat liver plasma membranes. The Biochemical journal 187, 381–392, doi:10.1042/bj1870381 (1980).

42 Afzal, F. et al. Differential regulation of β2 -adrenoceptor-mediated inotropic and lusitropic response by PDE3 and PDE4 in failing and non-failing rat cardiac ventricle. Br J Pharmacol 162, 54–71, doi: 10.1111/j.1476-5381.2010.00890.x (2011).

43 Zhang, J. H., Chung, T. D. & Oldenburg, K. R. A Simple Statistical Parameter for Use in Evaluation and Validation of High Throughput Screening Assays. Journal of biomolecular screening 4, 67–73 (1999).

44 Costello-Boerrigter, L. C. et al. Secretion of prohormone of B-type natriuretic peptide, proBNP1-108, is increased in heart failure. JACC Heart Fail 1, 207–212, doi:10.1016/j.jchf.2013.03.001 (2013).

45 Dickey, D. M. & Potter, L. R. ProBNP(1-108) is resistant to degradation and activates guanylyl cyclase-A with reduced potency. Clin Chem 57, 1272–1278, doi:10.1373/clinchem.2011.169151 (2011).

46 Antos, L. K. & Potter, L. R. Adenine nucleotides decrease the apparent Km of endogenous natriuretic peptide receptors for GTP. American journal of physiology. Endocrinology and metabolism 293, E1756–1763, doi: 10.1152/ajpendo.00321.2007 (2007).

47 Yoder, A. R., Stone, M. D., Griffin, T. J. & Potter, L. R. Mass spectrometric identification of phosphorylation sites in guanylyl cyclase A and B. Biochemistry 49, 10137–10145, doi:10.1021/bi101700e (2010).

48 Otto, N. M., McDowell, W. G., Dickey, D. M. & Potter, L. R. A Glutamate-Substituted Mutant Mimics the Phosphorylated and Active Form of Guanylyl Cyclase-A. Molecular Pharmacology 92, 67–74, doi:10.1124/mol.116.107995 (2017).

49 Kumar, R., Grammatikakis, N. & Chinkers, M. Regulation of the Atrial Natriuretic Peptide Receptor by Heat Shock Protein 90 Complexes*. Journal of Biological Chemistry 276, 11371–11375, doi:https://doi.org/10.1074/jbc.M010480200 (2001).

50 Jumper, J. et al. Highly accurate protein structure prediction with AlphaFold. Nature, doi:10.1038/s41586-021-03819-2 (2021).

51 Wenthur, C. J., Gentry, P. R., Mathews, T. P. & Lindsley, C. W. Drugs for allosteric sites on receptors. Annual review of pharmacology and toxicology 54, 165–184, doi:10.1146/annurev-pharmtox-010611-134525 (2014).

52 Lohse, M. J., Benovic, J. L., Caron, M. G. & Lefkowitz, R. J. Multiple pathways of rapid beta 2-adrenergic receptor desensitization. Delineation with specific inhibitors. Journal of Biological Chemistry 265, 3202–3209b, doi:https://doi.org/10.1016/S0021-9258(19)39754-6 (1990).

53 Neubig, R. R., Spedding, M., Kenakin, T. & Christopoulos, A. International Union of Pharmacology Committee on Receptor Nomenclature and Drug Classification. XXXVIII. Update on Terms and Symbols in Quantitative Pharmacology. Pharmacological Reviews 55, 597–606, doi:10.1124/pr.55.4.4 (2003).

