## Supplementary information for "Novel enhancers of guanylyl cyclase-A activity via allosteric modulation"

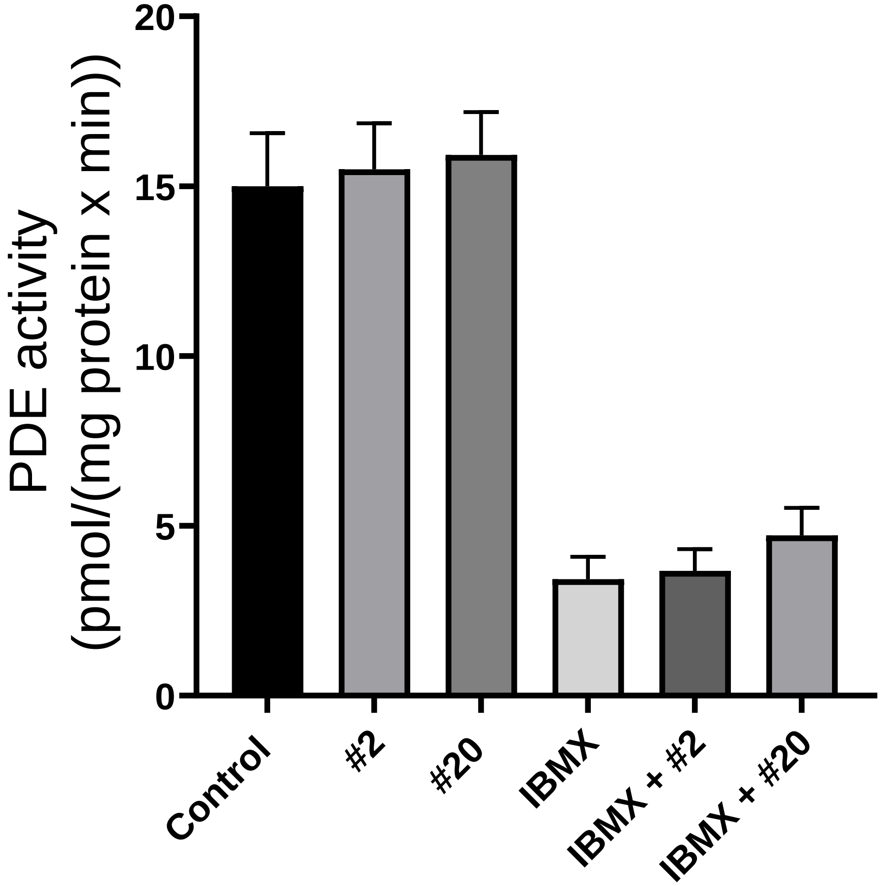


**Supplementary Fig. S1**

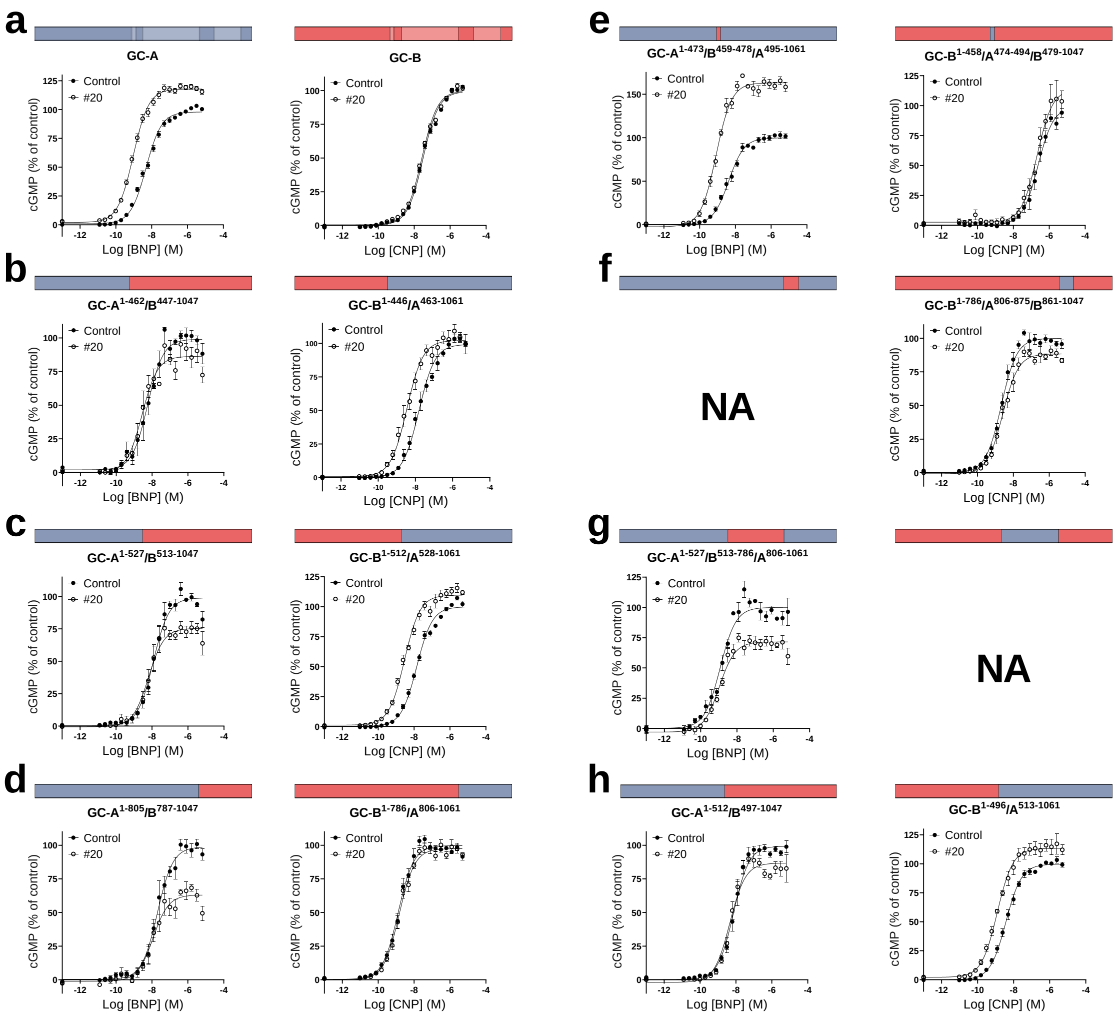

a
GC-A
GC-B
b
$GC-A^{1-462}/B^{447-1047}$
$GC-B^{1-446}/A^{463-1061}$
c
$GC-A^{1-527}/B^{513-1047}$
$GC-B^{1-512}/A^{528-1061}$
d
$GC-A^{1-805}/B^{787-1047}$
$GC-B^{1-786}/A^{806-1061}$
e
$GC-A^{1-473}/B^{459-478}/A^{495-1061}$
$GC-B^{1-458}/A^{474-494}/B^{479-1047}$
f
NA
$GC-B^{1-786}/A^{806-875}/B^{861-1047}$
g
$GC-A^{1-527}/B^{513-786}/A^{806-1061}$
NA
h
$GC-A^{1-512}/B^{497-1047}$
$GC-B^{1-496}/A^{513-1061}$
Control
#20
cGMP (% of control)
Log [BNP] (M)
Log [CNP] (M)

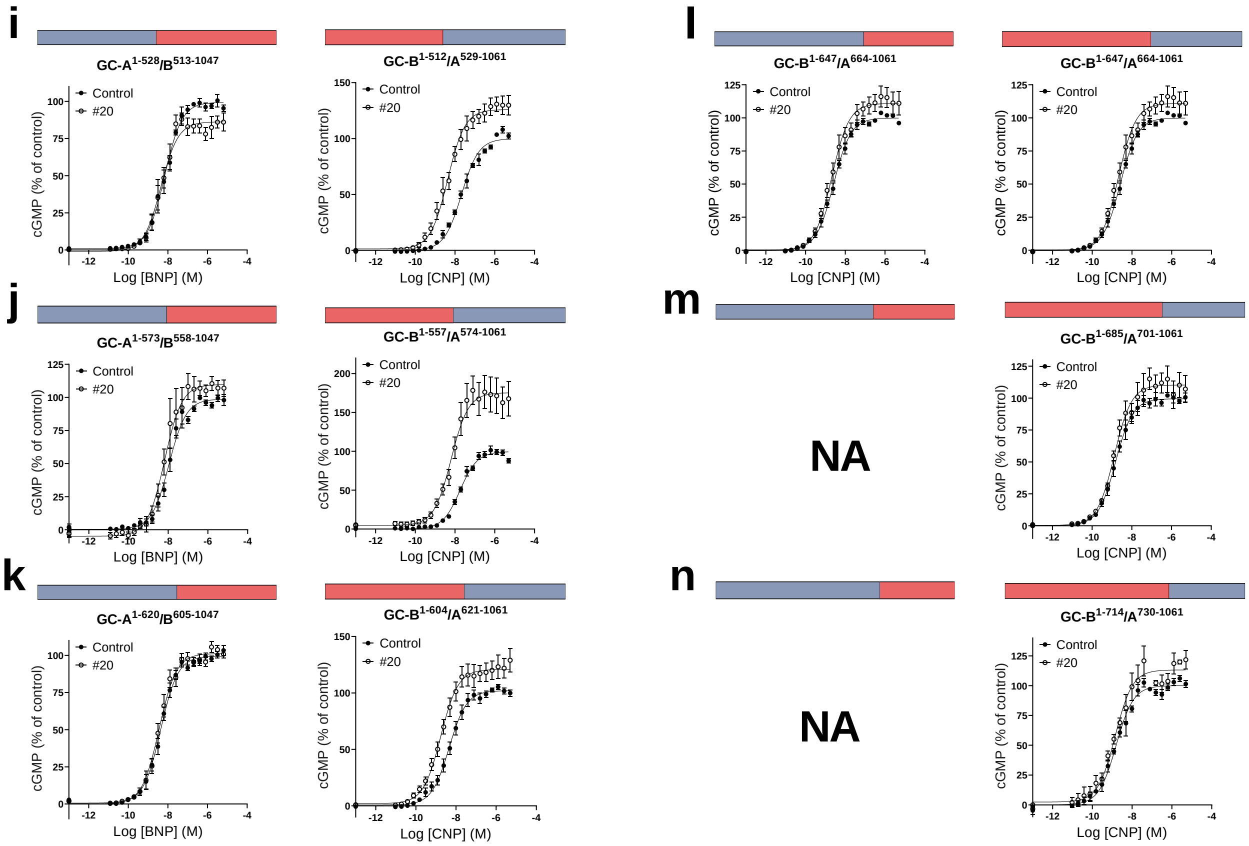


**Supplementary Fig. S2**

**GC-A** 621 QDILEN**E**SI**T**LDWMFRYSL**T**NDIVKGM**L**FLHN**GA**I**C**SHG**N**LKS 663
**GC-B** 605 QDILEN**D**SI**N**LDWMFRYSL**I**NDLVKGM**A**FLHN**SI**I**S**SHG**S**LKS 647

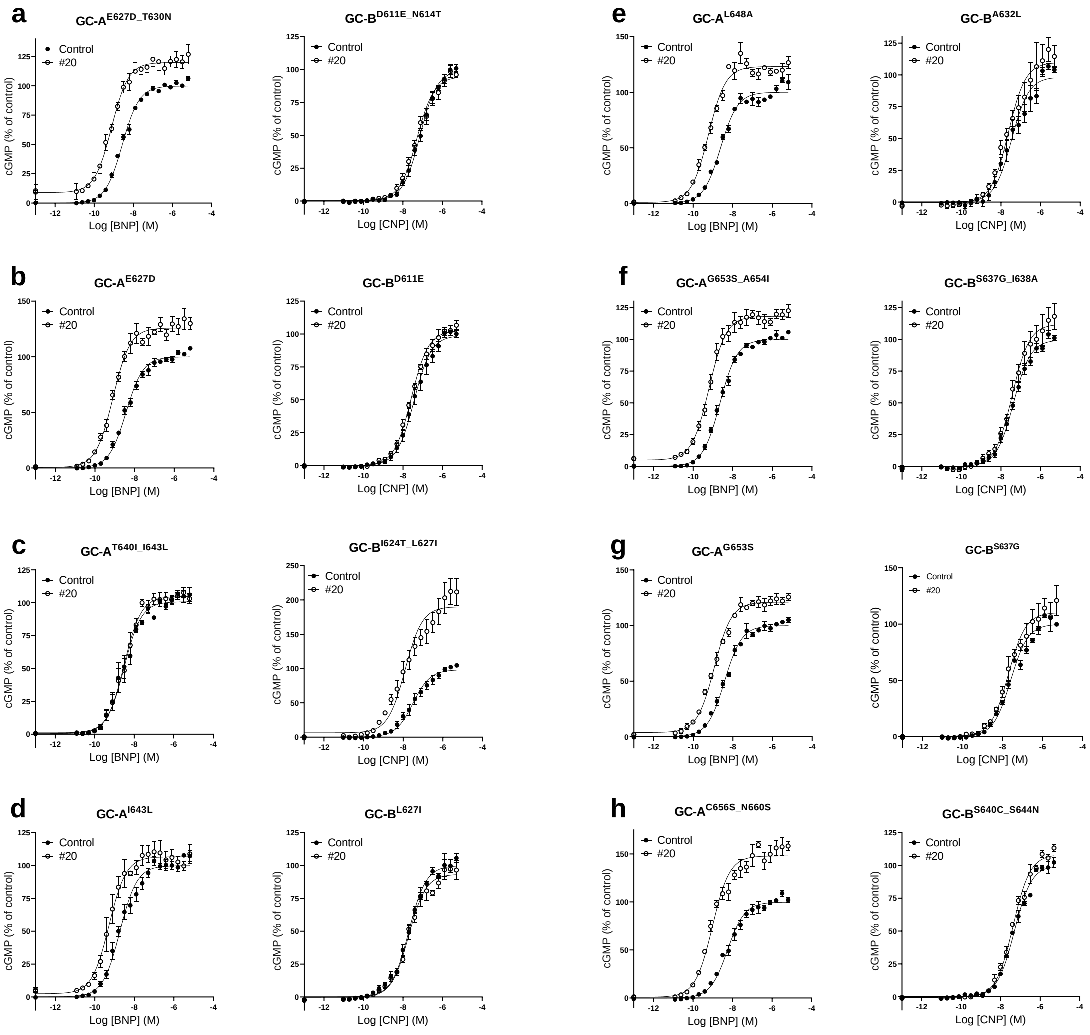


**Supplementary Fig. S3**

**Supplementary Table 1. a. Overview of primers used in constructing chimeric GC-A/B.** The pcDNA3.1(+) vector was linearized by restriction enzymes HindIII and XbaI and isolated from hGC-A or hGC-B pcDNA3.1(+) plasmids. Two or three DNA fragments were fused together with the linearized vector using the In-Fusion HD Enzyme premix that recognize a 15-20 bp overlap in the ends of each fragment. This overlap were added to the PCR primers.

| Chimeric receptor | Primers for GC-A fragment(s) | Primers for GC-B fragment(s) |
|---|---|---|
| GC-A$^{1-462}$/B$^{447-1047}$ | Forward: GCTAGCGTTTAAACTTAAGCTTACCATGCCCGGCCCCCGTAG<br>Reverse: GGTTTTGTCACAGCTAGGATCCTCATTGTCGAACCCAC | Forward: GGTTCGACAATGAGGATCCTAGCTGTGACAAAACCCCTCTGTC<br>Reverse: CAGCGGGTTTAAACGGGCCCTCTAGACTCGAGTTACAGCAGGCCG |
| GC-B$^{1-446}$/A$^{463-1061}$ | Forward: CATTCGATCTCGACGACCCCGCTTGCAACCAGGATCATCT<br>Reverse: GGTTTAAACGGGCCCTCTAGACTATCCGCGCGTGCTACTG | Forward: TAGCGTTTAAACTTAAGCTTACCATGGCCCTGCCGTCTTT<br>Reverse: AGATGATCCTGGTTGCAAGCGGGGTCGTCGAGATCGAATG |
| GC-A$^{1-494}$/B$^{479-1047}$ | Forward: GCTAGCGTTTAAACTTAAGCTTACCATGCCCGGCCCCCGTAG<br>Reverse: TTTTCGAGCATAAGTTTTCGGTAAATAAAGAAACTGACGA | Forward: TCGTCAGTTTCTTTATTTACCGAAAACTTATGCTCGAAAA<br>Reverse: CAGCGGGTTTAAACGGGCCCTCTAGACTCGAGTTACAGCAGGCCG |
| GC-B$^{1-478}$/A$^{495-1061}$ | Forward: TCAGCTCCTTCCTGATTTTTCGCAAGATGCAGCTGGAGAA<br>Reverse: GGTTTAAACGGGCCCTCTAGACTATCCGCGCGTGCTACTG | Forward: TAGCGTTTAAACTTAAGCTTACCATGGCCCTGCCGTCTTT<br>Reverse: TTCTCCAGCTGCATCTTGCGAAAAATCAGGAAGGAGCTGA |
| GC-A$^{1-527}$/B$^{513-1047}$ | Forward: GCTAGCGTTTAAACTTAAGCTTACCATGCCCGGCCCCCGTAG<br>Reverse: AGGGACAGAGTGAGCCGAGAGGCAGATCTCAGATGCCGCT | Forward: AGCGGCATCTGAGATCTGCCTCTCGGCTCACTCTGTCCCT<br>Reverse: CAGCGGGTTTAAACGGGCCCTCTAGACTCGAGTTACAGCAGGCCG |
| GC-B$^{1-512}$/A$^{528-1061}$ | Forward: GCTATCATAAGGGAGCCGGAGGGAGCCGCCTGACTCTCTC<br>Reverse: GGTTTAAACGGGCCCTCTAGACTATCCGCGCGTGCTACTG | Forward: TAGCGTTTAAACTTAAGCTTACCATGGCCCTGCCGTCTTT<br>Reverse: GAGAGAGTCAGGCGGCTCCCTCCGGCTCCCTTATGATAGC |
| GC-A$^{1-805}$/B$^{787-1047}$ | Forward: GCTAGCGTTTAAACTTAAGCTTACCATGCCCGGCCCCCGTAG<br>Reverse: CCCTCCTTATTAAAACGGCGGTTGAACTTCCTCAGGGTCA | Forward: TGACCCTGAGGAAGTTCAACCGCCGTTTTAATAAGGAGGG<br>Reverse: CAGCGGGTTTAAACGGGCCCTCTAGACTCGAGTTACAGCAGGCCG |
| GC-B$^{1-786}$/A$^{806-1061}$ | Forward: GGCAGATCAAGGGTTTTATTCGGGAGAATTCCTCTAATAT<br>Reverse: GGTTTAAACGGGCCCTCTAGACTATCCGCGCGTGCTACTG | Forward: TAGCGTTTAAACTTAAGCTTACCATGGCCCTGCCGTCTTT<br>Reverse: ATATTAGAGGAATTCTCCCGAATAAAACCCTTGATCTGCC |
| GC-A$^{1-875}$/B$^{861-1047}$ | Forward: GCTAGCGTTTAAACTTAAGCTTACCATGCCCGGCCCCCGTAG<br>Reverse: ATATCGGAGAAGTAGATGGTGACTGAGTCAAAGGCTTCGG | Forward: CCGAAGCCTTTGACTCAGTCACCATCTACTTCTCCGATAT<br>Reverse: CAGCGGGTTTAAACGGGCCCTCTAGACTCGAGTTACAGCAGGCCG |
| GC-B$^{1-860}$/A$^{876-1061}$ | Forward: CTGAGGCCTTTGACTCAGTGACTATCTATTTTTCCGACAT<br>Reverse: GGTTTAAACGGGCCCTCTAGACTATCCGCGCGTGCTACTG | Forward: TAGCGTTTAAACTTAAGCTTACCATGGCCCTGCCGTCTTT<br>Reverse: ATGTCGGAAAAATAGATAGTCACTGAGTCAAAGGCCTCAG |
| GC-A$^{1-462}$/B$^{447-458}$/A$^{474-1061}$ | 1. Forward: GCTAGCGTTTAAACTTAAGCTTACCATGCCCGGCCCCCGTAG<br>Reverse: AGAGGGGTTTTGTCACAGCTAGGATCCTCATTGTCGAACC<br>2. Forward: CTCTGTCCACACTCGCCATCGTGCTTGCACTGGTTGGCTC<br>Reverse: GGTTTAAACGGGCCCTCTAGACTATCCGCGCGTGCTACTG | Forward: GGTTCGACAATGAGGATCCTAGCTGTGACAAAACCCCTCT<br>Reverse: GAGCCAACCAGTGCAAGCACGATGGCGAGTGTGGACAGAG |
| GC-B$^{1-446}$/A$^{463-473}$/B$^{459-1047}$ | Forward: CATTCGATCTCGACGACCCCGCTTGCAACCAGGATCATCT<br>Reverse: ATGCCGGTGCCGAGTGCGACTTCCAAGGTACTAAGATGAT | 1. Forward: TAGCGTTTAAACTTAAGCTTACCATGGCCCTGCCGTCTTT<br>Reverse: AGATGATCCTGGTTGCAAGCGGGGTCGTCGAGATCGAATG<br>2. Forward: ATCATCTTAGTACCTTGGAAGTCGCACTCGGCACCGGCAT<br>Reverse: CAGCGGGTTTAAACGGGCCCTCTAGACTCGAGTTACAGCAGGCCG |
| GC-A$^{1-473}$/B$^{459-478}$/A$^{495-1061}$ | 1. Forward: GCTAGCGTTTAAACTTAAGCTTACCATGCCCGGCCCCCGTAG<br>Reverse: ATGCCGGTGCCGAGTGCGACTTCCAAGGTACTAAGATGAT<br>2. Forward: TCAGCTCCTTCCTGATTTTTCGCAAGATGCAGCTGGAGAA<br>Reverse: GGTTTAAACGGGCCCTCTAGACTATCCGCGCGTGCTACTG | Forward: ATCATCTTAGTACCTTGGAAGTCGCACTCGGCACCGGCAT<br>Reverse: TTCTCCAGCTGCATCTTGCGAAAAATCAGGAAGGAGCTGA |
| GC-B$^{1-458}$/A$^{474-494}$/B$^{479-1047}$ | Forward: CTCTGTCCACACTCGCCATCGTGCTTGCACTGGTTGGCTC<br>Reverse: TTTTCGAGCATAAGTTTTCGGTAAATAAAGAAACTGACGA | 1. Forward: TAGCGTTTAAACTTAAGCTTACCATGGCCCTGCCGTCTTT<br>Reverse: GAGCCAACCAGTGCAAGCACGATGGCGAGTGTGGACAGAG<br>2. Forward: TCGTCAGTTTCTTTATTTACCGAAAACTTATGCTCGAAAA<br>Reverse: CAGCGGGTTTAAACGGGCCCTCTAGACTCGAGTTACAGCAGGCCG |
| GC-A$^{1-527}$/B$^{513-786}$/A$^{806-1061}$ | 1. Forward: GCTAGCGTTTAAACTTAAGCTTACCATGCCCGGCCCCCGTAG<br>Reverse: AGGGACAGAGTGAGCCGAGA GGCAGATCTCAGATGCCGCT<br>2. Forward: GGCAGATCAAGGGTTTTATTCGGGAGAATTCCTCTAATAT<br>Reverse: GGTTTAAACGGGCCCTCTAGACTATCCGCGCGTGCTACTG | Forward: AGCGGCATCTGAGATCTGCCTCTCGGCTCACTCTGTCCCT<br>Reverse: ATATTAGAGGAATTCTCCCGAATAAAACCCTTGATCTGCC |

| | | |
|---|---|---|
| GC-B$^{1-512}$/A$^{528-805}$/B$^{787-1047}$ | Forward: GCTATCATAAGGGAGCCGGAGGGAGCCGCCTGACTCTCTC<br>Reverse: CCCTCCTTATTAAAACGGCGGTTGAACTTCCTCAGGGTCA | 1. Forward: TAGCGTTTAAACTTAAGCTTACCATGGCCCTGCCGTCTTT<br>Reverse: GAGAGAGTCAGGCGGCTCCCTCCGGCTCCCTTATGATAGC<br>2. Forward: TGACCCTGAGGAAGTTCAACCGCCGTTTTAATAAGGAGGG<br>Reverse: CAGCGGGTTTAAACGGGCCCTCTAGACTCGAGTTACAGCAGGCCG |
| GC-A$^{1-805}$/B$^{787-860}$/A$^{876-1061}$ | 1. Forward: GCTAGCGTTTAAACTTAAGCTTACCATGCCCGGCCCCCGTAG<br>Reverse: CCCTCCTTATTAAAACGGCGGTTGAACTTCCTCAGGGTCA<br>2. Forward: CTGAGGCCTTTGACTCAGTGACTATCTATTTTTCCGACAT<br>Reverse: GGTTTAAACGGGCCCTCTAGACTATCCGCGCGTGCTACTG | Forward: TGACCCTGAGGAAGTTCAACCGCCGTTTTAATAAGGAGGG<br>Reverse: ATGTCGGAAAAATAGATAGTCACTGAGTCAAAGGCCTCAG |
| GC-B$^{1-786}$/A$^{806-875}$/B$^{861-1047}$ | Forward: GGCAGATCAAGGGTTTTATTCGGGAGAATTCCTCTAATAT<br>Reverse: ATATCGGAGAAGTAGATGGTGACTGAGTCAAAGGCTTCGG | 1. Forward: TAGCGTTTAAACTTAAGCTTACCATGGCCCTGCCGTCTTT<br>Reverse: ATATTAGAGGAATTCTCCCGAATAAAACCCTTGATCTGCC<br>2. Forward: CCGAAGCCTTTGACTCAGTCACCATCTACTTCTCCGATAT<br>Reverse: CAGCGGGTTTAAACGGGCCCTCTAGACTCGAGTTACAGCAGGCCG |
| GC-A$^{1-527}$/B$^{513-860}$/A$^{876-1061}$ | 1. Forward: GCTAGCGTTTAAACTTAAGCTTACCATGCCCGGCCCCCGTAG<br>Reverse: AGGGACAGAGTGAGCCGAGAGGCAGATCTCAGATGCCGCT<br>2. Forward: CTGAGGCCTTTGACTCAGTGACTATCTATTTTTCCGACAT<br>Reverse: GGTTTAAACGGGCCCTCTAGACTATCCGCGCGTGCTACTG | Forward: AGCGGCATCTGAGATCTGCCTCTCGGCTCACTCTGTCCCT<br>Reverse: ATGTCGGAAAAATAGATAGTCACTGAGTCAAAGGCCTCAG |
| GC-B$^{1-512}$/A$^{528-875}$/B$^{861-1047}$ | Forward: GCTATCATAAGGGAGCCGGAGGGAGCCGCCTGACTCTCTC<br>Reverse: ATATCGGAGAAGTAGATGGTGACTGAGTCAAAGGCTTCGG | 1. Forward: TAGCGTTTAAACTTAAGCTTACCATGGCCCTGCCGTCTTT<br>Reverse: GAGAGAGTCAGGCGGCTCCCTCCGGCTCCCTTATGATAGC<br>2. Forward: CCGAAGCCTTTGACTCAGTCACCATCTACTTCTCCGATAT<br>Reverse: CAGCGGGTTTAAACGGGCCCTCTAGACTCGAGTTACAGCAGGCCG |
| GC-A$^{1-512}$/B$^{497-1047}$ | Forward: GCTAGCGTTTAAACTTAAGCTTACCATGCCCGGCCCCCGTAG<br>Reverse: TTTCCGAACTGCAATTCCTCCCACCTAACACGCCACAGCT | Forward: AGCTGTGGCGTGTTAGGTGGGAGGAATTGCAGTTCGGAAA<br>Reverse: CAGCGGGTTTAAACGGGCCCTCTAGACTCGAGTTACAGCAGGCCG |
| GC-B$^{1-496}$/A$^{513-1061}$ | Forward: TGCTTTGGCGAATCAGGTGGGAGGATGTGGAACCATCAAG<br>Reverse: GGTTTAAACGGGCCCTCTAGACTATCCGCGCGTGCTACTG | Forward: TAGCGTTTAAACTTAAGCTTACCATGGCCCTGCCGTCTTT<br>Reverse: CTTGATGGTTCCACATCCTCCCACCTGATTCGCCAAAGCA |
| GC-A$^{1-528}$/B$^{513-1047}$ | Forward: GCTAGCGTTTAAACTTAAGCTTACCATGCCCGGCCCCCGTAG<br>Reverse: AGGGACAGAGTGAGCCGAGACCCGGCAGATCTCAGATGCC | Forward: GGCATCTGAGATCTGCCGGGTCTCGGCTCACTCTGTCCCT<br>Reverse: CAGCGGGTTTAAACGGGCCCTCTAGACTCGAGTTACAGCAGGCCG |
| GC-B$^{1-512}$/A$^{529-1061}$ | Forward: GCTATCATAAGGGAGCCGGAAGCCGCCTGACTCTCTCCGG<br>Reverse: GGTTTAAACGGGCCCTCTAGACTATCCGCGCGTGCTACTG | Forward: TAGCGTTTAAACTTAAGCTTACCATGGCCCTGCCGTCTTT<br>Reverse: CCGGAGAGAGTCAGGCGGCTTCCGGCTCCCTTATGATAGC |
| GC-A$^{1-573}$/B$^{558-1047}$ | Forward: GCTAGCGTTTAAACTTAAGCTTACCATGCCCGGCCCCCGTAG<br>Reverse: ACCTGACGAGTAAGCTCGATCCGCTTGCGGTTCACCCGCT | Forward: AGCGGGTGAACCGCAAGCGGATCGAGCTTACTCGTCAGGT<br>Reverse: CAGCGGGTTTAAACGGGCCCTCTAGACTCGAGTTACAGCAGGCCG |
| GC-B$^{1-557}$/A$^{574-1061}$ | Forward: AGCACGTGAATAAGAAGCGGATTGAGCTGACGCGCAAAGT<br>Reverse: GGTTTAAACGGGCCCTCTAGACTATCCGCGCGTGCTACTG | Forward: TAGCGTTTAAACTTAAGCTTACCATGGCCCTGCCGTCTTT<br>Reverse: ACTTTGCGCGTCAGCTCAATCCGCTTCTTATTCACGTGCT |
| GC-A$^{1-620}$/B$^{605-1047}$ | Forward: GCTAGCGTTTAAACTTAAGCTTACCATGCCCGGCCCCCGTAG<br>Reverse: TCGTTTTCCAGAATATCCTGGAGAGAGCCCCGTGGGCAAT | Forward: ATTGCCCACGGGGCTCTCTCCAGGATATTCTGGAAAACGA<br>Reverse: CAGCGGGTTTAAACGGGCCCTCTAGACTCGAGTTACAGCAGGCCG |
| GC-B$^{1-604}$/A$^{621-1061}$ | Forward: ATTGTCCGCGGGGAAGTTTGCAGGACATTCTCGAGAACGA<br>Reverse: GGTTTAAACGGGCCCTCTAGACTATCCGCGCGTGCTACTG | Forward: TAGCGTTTAAACTTAAGCTTACCATGGCCCTGCCGTCTTT<br>Reverse: TCGTTCTCGAGAATGTCCTGCAAACTTCCCCGCGGACAAT |
| GC-A$^{1-663}$/B$^{648-1047}$ | Forward: GCTAGCGTTTAAACTTAAGCTTACCATGCCCGGCCCCCGTAG<br>Reverse: CTGTCAACCACACAGTTAGATGACTTAAGGTTTCCGTGGG | Forward: CCCACGGAAACCTTAAGTCATCTAACTGTGTGGTTGACAG<br>Reverse: CAGCGGGTTTAAACGGGCCCTCTAGACTCGAGTTACAGCAGGCCG |
| GC-B$^{1-647}$/A$^{664-1061}$ | Forward: CACACGGATCACTTAAAAGCTCAAACTGTGTGGTGGACGG<br>Reverse: GGTTTAAACGGGCCCTCTAGACTATCCGCGCGTGCTACTG | Forward: TAGCGTTTAAACTTAAGCTTACCATGGCCCTGCCGTCTTT<br>Reverse: CCGTCCACCACACAGTTTGAGCTTTTAAGTGATCCGTGTG |
| GC-A$^{1-700}$/B$^{686-1047}$ | Forward: GCTAGCGTTTAAACTTAAGCTTACCATGCCCGGCCCCCGTAG<br>Reverse: AGCAACTCCGGGGCGGTCCACAGCTTCTTTGCGTAGACGG | Forward: CCGTCTACGCAAAGAAGCTGTGGACCGCCCCGGAGTTGCT<br>Reverse: CAGCGGGTTTAAACGGGCCCTCTAGACTCGAGTTACAGCAGGCCG |
| GC-B$^{1-685}$/A$^{701-1061}$ | Forward: CCCTGTACGCCAAGAAGCTGTGGACCGCTCCCGAGCTGCT<br>Reverse: GGTTTAAACGGGCCCTCTAGACTATCCGCGCGTGCTACTG | Forward: TAGCGTTTAAACTTAAGCTTACCATGGCCCTGCCGTCTTT<br>Reverse: AGCAGCTCGGGAGCGGTCCACAGCTTCTTGGCGTACAGGG |
| GC-A$^{1-729}$/B$^{715-1047}$ | Forward: GCTAGCGTTTAAACTTAAGCTTACCATGCCCGGCCCCCGTAG<br>Reverse: GAGCGCAGAGCAATTTCCTGCAGTATGATCCCGAAGGAGT | Forward: ACTCCTTCGGGATCATACTGCAGGAAATTGCTCTGCGCTC<br>Reverse: CAGCGGGTTTAAACGGGCCCTCTAGACTCGAGTTACAGCAGGCCG |
| GC-B$^{1-714}$/A$^{730-1061}$ | Forward: ACTCCTTTGGAATTATCCTGCAGGAAATTGCTCTTCGGAG<br>Reverse: GGTTTAAACGGGCCCTCTAGACTATCCGCGCGTGCTACTG | Forward: TAGCGTTTAAACTTAAGCTTACCATGGCCCTGCCGTCTTT<br>Reverse: CTCCGAAGAGCAATTTCCTGCAGGATAATTCCAAAGGAGT |

| | | |
|---|---|---|
| GC-$A^{1-620}/B^{605-714}/A^{730-1061}$ | 1. Forward: GCTAGCGTTTAAACTTAAGCTTACCATGCCCGGCCCCCGTAG<br>Reverse: TCGTTTTCCAGAATATCCTGGAGAGAGCCCCGTGGGCAAT<br>2. Forward: ACTCCTTTGGAATTATCCTGCAGGAAATTGCTCTTCGGAG<br>Reverse: GGTTTAAACGGGCCCTCTAGACTATCCGCGCGTGCTACTG | Forward: ATTGCCCACGGGGCTCTCTCCAGGATATTCTGGAAAACGA<br>Reverse: CTCCGAAGAGCAATTTCCTGCAGGATAATTCCAAAGGAGT |
| GC-$B^{1-604}/A^{621-729}/B^{715-1047}$ | Forward: ATTGTCCGCGGGGAAGTTTGCAGGACATTCTCGAGAACGA<br>Reverse: GAGCGCAGAGCAATTTCCTGCAGTATGATCCCGAAGGAGT | 1. Forward: TAGCGTTTAAACTTAAGCTTACCATGGCCCTGCCGTCTTT<br>Reverse: TCGTTCTCGAGAATGTCCTGCAAACTTCCCCGCGGACAAT<br>2. Forward: ACTCCTTCGGGATCATACTGCAGGAAATTGCTCTGCGCTC<br>Reverse: CAGCGGGTTTAAACGGGCCCTCTAGACTCGAGTTACAGCAGGCCG |
| GC-$A^{1-620}/B^{605-647}/A^{664-1061}$ | 1. Forward: GCTAGCGTTTAAACTTAAGCTTACCATGCCCGGCCCCCGTAG<br>Reverse: TCGTTTTCCAGAATATCCTGGAGAGAGCCCCGTGGGCAAT<br>2. Forward: CACACGGATCACTTAAAAGCTCAAACTGTGTGGTGGACGG<br>Reverse: GGTTTAAACGGGCCCTCTAGACTATCCGCGCGTGCTACTG | Forward: ATTGCCCACGGGGCTCTCTCCAGGATATTCTGGAAAACGA<br>Reverse: CCGTCCACCACACAGTTTGAGCTTTTAAGTGATCCGTGTG |
| GC-$B^{1-604}/A^{621-663}/B^{648-1047}$ | Forward: ATTGTCCGCGGGGAAGTTTGCAGGACATTCTCGAGAACGA<br>Reverse: CTGTCAACCACACAGTTAGATGACTTAAGGTTTCCGTGGG | 1. Forward: TAGCGTTTAAACTTAAGCTTACCATGGCCCTGCCGTCTTT<br>Reverse: TCGTTCTCGAGAATGTCCTGCAAACTTCCCCGCGGACAAT<br>2. Forward: CCCACGGAAACCTTAAGTCATCTAACTGTGTGGTTGACAG<br>Reverse: CAGCGGGTTTAAACGGGCCCTCTAGACTCGAGTTACAGCAGGCCG |
| GC-$A^{1-663}/B^{648-686}/A^{701-1061}$ | 1. Forward: GCTAGCGTTTAAACTTAAGCTTACCATGCCCGGCCCCCGTAG<br>Reverse: CTGTCAACCACACAGTTAGATGACTTAAGGTTTCCGTGGG<br>2. Forward: CCCTGTACGCCAAGAAGCTGTGGACCGCTCCCGAGCTGCT<br>Reverse: GGTTTAAACGGGCCCTCTAGACTATCCGCGCGTGCTACTG | Forward: CCCACGGAAACCTTAAGTCATCTAACTGTGTGGTTGACAG<br>Reverse: AGCAGCTCGGGAGCGGTCCACAGCTTCTTGGCGTACAGGG |
| GC-$B^{1-647}/A^{664-700}/B^{686-1047}$ | Forward: CACACGGATCACTTAAAAGCTCAAACTGTGTGGTGGACGG<br>Reverse: AGCAACTCCGGGGCGGTCCACAGCTTCTTTGCGTAGACGG | 1. Forward: TAGCGTTTAAACTTAAGCTTACCATGGCCCTGCCGTCTTT<br>Reverse: CCGTCCACCACACAGTTTGAGCTTTTAAGTGATCCGTGTG<br>2. Forward: CCGTCTACGCAAAGAAGCTGTGGACCGCCCCGGAGTTGCT<br>Reverse: CAGCGGGTTTAAACGGGCCCTCTAGACTCGAGTTACAGCAGGCCG |
| GC-$A^{1-700}/B^{686-715}/A^{731-1061}$ | 1. Forward: GCTAGCGTTTAAACTTAAGCTTACCATGCCCGGCCCCCGTAG<br>Reverse: AGCAACTCCGGGGCGGTCCACAGCTTCTTTGCGTAGACGG<br>2. Forward: ACTCCTTTGGAATTATCCTGCAGGAAATTGCTCTTCGGAG<br>Reverse: GGTTTAAACGGGCCCTCTAGACTATCCGCGCGTGCTACTG | Forward: CCGTCTACGCAAAGAAGCTGTGGACCGCCCCGGAGTTGCT<br>Reverse: CTCCGAAGAGCAATTTCCTGCAGGATAATTCCAAAGGAGT |
| GC-$B^{1-685}/A^{701-730}/B^{716-1047}$ | Forward: CCCTGTACGCCAAGAAGCTGTGGACCGCTCCCGAGCTGCT<br>Reverse: GAGCGCAGAGCAATTTCCTGCAGTATGATCCCGAAGGAGT | 1. Forward: TAGCGTTTAAACTTAAGCTTACCATGGCCCTGCCGTCTTT<br>Reverse: AGCAGCTCGGGAGCGGTCCACAGCTTCTTGGCGTACAGGG<br>2. Forward: ACTCCTTCGGGATCATACTGCAGGAAATTGCTCTGCGCTC<br>Reverse: CAGCGGGTTTAAACGGGCCCTCTAGACTCGAGTTACAGCAGGCCG |

**b. Overview of primers used in constructing single or dual mutations in GC-A and GC-B.** Primers for mutagenesis were mainly design using the web based In-Fusion Cloning Primer Design Tool from Takara Bio (available from https://www.takarabio.com/learning-centers/cloning/primer-design-and-other-tools). The primers have a 15 bp overlap in the 5' end and the mutation incorporated. GC-A or GC-B was used as templates unless otherwise is specified.

| Construct | Primers |
|---|---|
| GC-A$^{E627D}$ | Forward: CGAGAACGACTCCATTACACTGGACTGGATGTTCC<br>Reverse: ATGGAGTCGTTCTCGAGAATGTCCTGGAGAG |
| GC-A$^{E627D_T630N}$ | Using GC-A$^{E627D}$ as template:<br>Forward: CTCCATTAACCTGGACTGGATGTTCCGC<br>Reverse: TCCAGGTTAATGGAGTCGTTCTCGAGAATGTC |
| GC-A$^{T640I}$ | Forward: TAGTCTGATTAATGACATCGTCAAGGGGATGC<br>Reverse: TCATTAATCAGACTATAGCGGAACATCCAGTCC |
| GC-A$^{I643L}$ | Forward: TAATGACCTTGTCAAGGGGATGCTCTTCC<br>Reverse: TTGACAAGGTCATTAGTCAGACTATAGCGGAAC |
| GC-A$^{T640I_I643L}$ | Forward: TAGTCTGATTAATGACCTTGTCAAGGGGATGCTCTTCC<br>Reverse: TTGACAAGGTCATTAATCAGACTATAGCGGAACATCCAGT |
| GC-A$^{L648A}$ | Forward: GGGGATGGCTTTCCTTCACAACGGTGCC<br>Reverse: AGGAAAGCCATCCCCTTGACGATGTCATTAGTC |
| GC-A$^{G653S}$ | Forward: TCACAACTCCGCCATCTGCTCCCACGGA<br>Reverse: ATGGCGGAGTTGTGAAGGAAGAGCATCCCCT |
| GC-A$^{G653S_A654I}$ | Using GC-A$^{G653S}$ as template:<br>Forward: CAACTCCATTATCTGCTCCCACGGAAACC<br>Reverse: CAGATAATGGAGTTGTGAAGGAAGAGCATCC |
| GC-A$^{C656S_N660S}$ | First constructed GC-A$^{C656S}$ using<br>Forward: TGCCATCTCATCCCACGGAAACCTTAAGTC<br>Reverse: TGGGATGAGATGGCACCGTTGTGAAGG<br>Then constructed GC-A$^{C656S_N660S}$ using GC-A$^{C656S}$ as the template and primers:<br>Forward: CCACGGATCACTTAAGTCATCAAACTGTGTGGTGG<br>Reverse: TTAAGTGATCCGTGGGATGAGATGGC |
| GC-B$^{D611E}$ | Forward: GGAAAACGAGAGCATCAACCTTGATTGGATGTTC<br>Reverse: ATGCTCTCGTTTTCCAGAATATCCTGCAAAC |
| GC-B$^{D611E_N614T}$ | Using GC-B$^{D611E}$ as template:<br>Forward: GAGCATCACACTTGATTGGATGTTCCGCTACTCC<br>Reverse: TCAAGTGTGATGCTCTCGTTTTCCAGAATATCC |
| GC-B$^{I624T}$ | Forward: CTCCCTGACTAACGACCTTGTGAAAGGTATGGCT<br>Reverse: TCGTTAGTCAGGGAGTAGCGGAACATCCAA |
| GC-B$^{L627I}$ | Forward: TAACGACATCGTGAAAGGTATGGCTTTCCTGC<br>Reverse: TTCACGATGTCGTTAATCAGGGAGTAGCGG |
| GC-B$^{I624T_L627I}$ | Forward: CTCCCTGACTAACGACATCGTGAAAGGTATGGCTTTCCTG<br>Reverse: TTCACGATGTCGTTAGTCAGGGAGTAGCGGAACATCCAA |
| GC-B$^{A632L}$ | Forward: AGGTATGCTGTTCCTGCACAATTCCATTATCTCA<br>Reverse: AGGAACAGCATACCTTTCACAAGGTCGTT |
| GC-B$^{S637G}$ | Forward: GCACAATGGTATTATCTCATCACACGGATCACT<br>Reverse: ATAATACCATTGTGCAGGAAAGCCATACCT |

| | |
|---|---|
| GC-B$^{S637G_I638A}$ | Using GC-B$^{S637G}$ as template:<br>Forward: CAATGGTGCCATCTCATCACACGGATCACTT<br>Reverse: GAGATGGCACCATTGTGCAGGAAAGCCA |
| GC-B$^{S640C_S644N}$ | First constructed GC-B$^{S640C}$ using<br>Forward: CATTATCTGCTCACACGGATCACTTAAAAGC<br>Reverse: TGTGAGCAGATAATGGAATTGTGCAGGAAAGC<br>Then constructed GC-B$^{S640C_S644N}$ using GC-B$^{S640C}$ as the template and primers:<br>Forward: ACACGGAAACCTTAAAAGCTCTAACTGTGTGGTTG<br>Reverse: TTAAGGTTTCCGTGTGAGCAGATAATGG |
| **GC-AT640 and GC-BI624 focused constructs** | |
| GC-A$^{T640A}$ | Forward: TAGTCTGGCTAATGACATCGTCAAGGGGATGC<br>Reverse: TCATTAGCCAGACTATAGCGGAACATCCAGTCC |
| GC-A$^{T640D}$ | Forward: TAGTCTGGATAATGACATCGTCAAGGGGATGC<br>Reverse: TCATTATCCAGACTATAGCGGAACATCCAGTCC |
| GC-A$^{T640E}$ | Forward: TAGTCTGGAGAATGACATCGTCAAGGGGATGC<br>Reverse: TCATTCTCCAGACTATAGCGGAACATCCAGTCC |
| GC-A$^{T640L}$ | Forward: TAGTCTGCTTAATGACATCGTCAAGGGGATGC<br>Reverse: TCATTAAGCAGACTATAGCGGAACATCCAGTCC |
| GC-A$^{T640S}$ | Forward: TAGTCTGAGTAATGACATCGTCAAGGGGATGC<br>Reverse: TCATTACTCAGACTATAGCGGAACATCCAGTCC |
| GC-A$^{T640Y}$ | Forward: TAGTCTGTACAATGACATCGTCAAGGGGATGC<br>Reverse: TCATTGTACAGACTATAGCGGAACATCCAGTCC |
| GC-A$^{T640V}$ | Forward: TAGTCTGGTTAATGACATCGTCAAGGGGATGC<br>Reverse: TCATTAACCAGACTATAGCGGAACATCCAGTCC |
| GC-B$^{I624A}$ | Forward: CTCCCTGGCTAACGACCTTGTGAAAGGTATGGCT<br>Reverse: TCGTTAGCCAGGGAGTAGCGGAACATCCAA |
| GC-B$^{I624D}$ | Forward: CTCCCTGGATAACGACCTTGTGAAAGGTATGGCT<br>Reverse: TCGTTATCCAGGGAGTAGCGGAACATCCAA |
| GC-B$^{I624L}$ | Forward: CTCCCTGCTTAACGACCTTGTGAAAGGTATGGCT<br>Reverse: TCGTTAAGCAGGGAGTAGCGGAACATCCAA |
| GC-B$^{I624S}$ | Forward: CTCCCTGAGTAACGACCTTGTGAAAGGTATGGCT<br>Reverse: TCGTTACTCAGGGAGTAGCGGAACATCCAA |
| GC-B$^{I624Y}$ | Forward: CTCCCTGTACAACGACCTTGTGAAAGGTATGGCT<br>Reverse: TCGTTGTACAGGGAGTAGCGGAACATCCAA |
| GC-B$^{I624V}$ | Forward: CTCCCTGGTTAACGACCTTGTGAAAGGTATGGCT<br>Reverse: TCGTTAACCAGGGAGTAGCGGAACATCCAA |
| **GC$^{7E}$** | |
| GC-A$^{S519E}$ | Forward: ACCATCAGAGCTGGAGCGGCATCTGAGATCTG<br>Reverse: TCCAGCTCTGATGGTTCCACATCCTCCCAC |
| GC-A$^{S519E_S529E}$ | Using GC-A$^{S519E}$ as template:<br>Forward: TGCCGGGGAACGCCTGACTCTCTCCGGAC<br>Reverse: AGGCGTTCCCCGGCAGATCTCAGATGC |
| GC-A$^{S519E_S529E_T532E}$ | Using GC-A$^{S519E_S529E}$ as template:<br>Forward: ACGCCTGGAACTCTCCGGACGAGGCTCCA<br>Reverse: GAGAGTTCCAGGCGTTCCCCGGCAGA |
| GC-A$^{S519E_S529E_T532E_S534E}$ | Using GC-A$^{S519E_S529E_T532E}$ as template:<br>Forward: GGAACTCGAAGGACGAGGCTCCAACTACGG<br>Reverse: CGTCCTTCGAGTTCCAGGCGTTCCCC |

| | |
|---|---|
| GC-A$^{\text{S519E_S529E_T532E_S534E_S538E}}$ | Using GC-A$^{\text{S519E_S529E_T532E_S534E}}$ as template:<br>Forward: ACGAGGCGAGAACTACGGTTCTTTGCTGACAACC<br>Reverse: TAGTTCTCGCCTCGTCCTTCGAGTTCC |
| GC-_A$^{\text{S519E_S529E_T532E_S534E_S538E_S542E}}$ | Using GC-A$^{\text{S519E_S529E_T532E_S534E_S538E}}$ as template:<br>Forward: CTACGGTGAGTTGCTGACAACCGAAGGCCA<br>Reverse: AGCAACTCACCGTAGTTCTCGCCTCGT |
| GC-A$^{\text{S519E_S529E_T532E_S534E_S538E_S542E_T545E}}$ | Using GC-_A$^{\text{S519E_S529E_T532E_S534E_S538E_S542E}}$ as template:<br>Forward: GTTGCTGGAAACCGAAGGCCAGTTCCAGG<br>Reverse: TCGGTTTCCAGCAACTCACCGTAGTTCTCG |

**FIGURE LEGENDS:**

Supplementary Fig. S1

**Compounds did not inhibit PDE activity.** cGMP-PDE activity in GC-A-expressing cells treated with 0.1% DMSO (control) or 10 µM compound #2 or #20 in the presence or absence of the general PDE inhibitor IBMX. Data points are means±SEM (n=3-7)

Supplementary Fig. S2

**All chimeric GC-A/B tested.** Concentration-response curves for chimeric GC-A/B pairs with the indicated concentrations of BNP (GC-A extracellular domain) or CNP (GC-B extracellular domain) stimulated with 0.1% DMSO (control) or 10 µM compound #20. Some chimeric receptors were not active (NA). Data points are means±SEM (n=3-6 for chimeric GC-A/B pairs, n=53 for GC-A and GC-B).

Supplementary Fig. S3

**Mutations of non-conserved amino acids reveal that the activity only occurred in the presence of GC-A$^{T640}$.** Concentration-response curves for BNP and CNP and the effects of compound #20 towards GC mutations with single or dual amino acid swapping of non-conserved amino acids. In the region 621-663 in GC-A, only nine amino acids are non-conserved between GC-A and GC-B. Data points are means±SEM (n=3-5).
